# *N*-acyl homoserine lactone signaling modulates bacterial community associated with human dental plaque

**DOI:** 10.1101/2024.03.15.585217

**Authors:** Rakesh Sikdar, Mai V. Beauclaire, Bruno P. Lima, Mark C. Herzberg, Mikael H. Elias

## Abstract

*N*-acyl homoserine lactones (AHLs) are small diffusible signaling molecules that mediate a cell density-dependent bacterial communication system known as quorum sensing (QS). AHL-mediated QS regulates gene expression to control many critical bacterial behaviors including biofilm formation, pathogenicity, and antimicrobial resistance. Dental plaque is a complex multispecies oral biofilm formed by successive colonization of the tooth surface by groups of commensal, symbiotic, and pathogenic bacteria, which can contribute to tooth decay and periodontal diseases. While the existence and roles of AHL-mediated QS in oral microbiota have been debated, recent evidence indicates that AHLs play significant roles in oral biofilm development and community dysbiosis. The underlying mechanisms, however, remain poorly characterized. To better understand the importance of AHL signaling in dental plaque formation, we manipulated AHL signaling by adding AHL lactonases or exogenous AHL signaling molecules. We find that AHLs can be detected in dental plaque grown under 5% CO_2_ conditions, but not when grown under anaerobic conditions, and yet anaerobic cultures are still responsive to AHLs. QS signal disruption using lactonases leads to changes in microbial population structures in both planktonic and biofilm states, changes that are dependent on the substrate preference of the used lactonase but mainly result in the increase in the abundance of commensal and pioneer colonizer species. Remarkably, the opposite manipulation, that is the addition of exogenous AHLs increases the abundance of late colonizer bacterial species. Hence, this work highlights the importance of AHL-mediated QS in dental plaque communities, its potential different roles in anaerobic and aerobic parts of dental plaque, and underscores the potential of QS interference in the control of periodontal diseases

## Introduction

Quorum sensing (QS) is a cell-density-dependent bacterial communication system mediated by small diffusible signaling molecules called autoinducers (AI) [1,2]. There are three major types of AIs:(i) AI-1 (*N*-acyl homoserine lactones or AHLs) [3–5]; (ii) AI-2 [6–9] consisting of 4,5-dihydroxy-2,3-pentandedione (DPD) derivatives [10]; and (iii) autoinducing peptides (AIPs), specific to *Gram*-positive bacteria [11]. AIs are typically secreted into the environment and can re-enter the cells either by passive diffusion in the case of AI-1 [3] or via active transport for AI-2 and AIPs [6,12,13]. AIs bind to their target receptors upon re-entry, regulating the expression of a wide variety of genes [2]. Indeed, QS can regulate large parts of the bacterial genome (up to 26%) [14]. By controlling gene expression, QS regulates several bacterial behaviors such as biofilm formation, virulence, and antimicrobial resistance [15,16]. These behaviors are critical for bacterial adaptation to varying ecological niches often characterized by hostile environmental conditions and inconsistent nutrient availability [17].

Dental plaque is a complex oral biofilm, composed of commensal, symbiotic, and pathogenic members of oral microbiota, adhering to the tooth surface [18]. The “core” microbiome of the human oral cavity is normally comprised of thousands of species of microorganisms including bacteria, fungi, archaea, protozoa, and viruses [19,20]. Organisms from these kingdoms inhabit distinct ecological niches found in unique environments that includes gingival sulcus, tongue, cheek, soft and hard palates, throat, saliva, and teeth [21,22]. The bacterial taxonomic diversity in the oral cavity includes about 700 species belonging to 185 genera and 12 phyla [23,24]. The healthy oral microbiome is dominated by Gram-positive commensal bacteria such as *Streptococcus* and *Actinomyces* ssp. with high propensity for fermentation. [25–27]. Abrupt ecological changes and alterations of host-associated innate and adaptive immunity in the oral cavity can cause dysbiosis. Dysbiosis is characterized by an increased prevalence of anaerobic Gram-negative pathogenic bacteria with high proteolytic activity such as *Porphyromonas* [28], often leading to the development of periodontal diseases [25–27]. Perturbations in oral communities can also favor the emergence of more acidogenic bacteria associated with dental caries [29].

Dental plaque formation is characterized by successive spatial and temporal colonization of the tooth surface requiring cooperative and competitive interactions between several groups of bacteria [30–32]. Interbacterial interactions are hypothesized to depend in part on QS-dependent regulation of cellular metabolism, physiology, secretion, virulence, motility, attachment, and cooperativity/competition among the members of the oral microbiota [33,34]. AI-2 and AIPs are known to be important in the development of oral biofilms. Many dental plaque colonizers were shown to produce and/or respond to AI-2, [35–42]. Gram-positive pioneer colonizers like oral *Streptococcus* species can utilize AIPs [43–46]. Hence, AI-2/AIP-mediated QS appears to be the primary mode of signaling and regulation that contributes to the formation and development of oral biofilms including dental plaque [34]. The potential role of AI-1/AHL-based QS in the development of dental plaque and other oral biofilms remains elusive. In fact, AHLs were long undetected in cultures of pathogenic oral microbiota [37,47,48]. Given the lack of identified AHL synthases and receptor homologs in plaque pathogens, AHLs have been considered to make an insignificant contribution to oral biofilm formation [49,50].

This paradigm may be ripe for change. Various types of AHLs are found in human saliva samples [51–53]. Sources include the AHL-producing Gram-negative bacteria *Pseudomonas putida* [54]*, Enterobacter* [55]*, Klebsiella pneumoniae* [56]*, Citrobacter amalonaticus* [57] *and Burkholderia* [58], found on tongue surfaces and within dentinal caries. Certain strains of the oral pathogen *Porphyromonas gingivalis* also possess AHL synthase/receptor homologs [59,60] and produce small quantities of AHLs in both axenic and multispecies cultures [53]. Indeed, putative AHL biosynthetic genes exist in oral microbial genomes [61]. Metagenomic analysis of human dental plaque revealed a high abundance of AHL-synthase (HdtS), receptor homologs and Quorum Quenching (QQ) enzymes [62]. Muras and colleagues have recently shown [51,53,62] that while exogenously added AHLs promote the growth of pathogens in oral communities, AHL-degrading QQ enzymes inhibit oral biofilm formation and alter their microbial composition.

Yet, the specific role(s) of AHL-mediated QS in oral communities remain unclear. To increase our understanding of the importance of AI-1, one strategy is to apply different AHL signals and AHL-degrading lactonases onto microbial communities and evaluate their effects. Lactonases are QQ enzymes that hydrolyze the lactone ring of AHLs and disrupt AHL-mediated QS [63–66]. We previously characterized a number of lactonases, including SsoPox from the Phosphotriesterase-like Lactonase (PLL) family [67–69] and GcL from the Metallo-β-Lactamase (MLL) family [70]. Interestingly, these two lactonases exhibit distinct, yet overlapping substrate specificities: SsoPox preferentially degrades long chain AHLs (C8 or longer) [69] and GcL is a generalist lactonase with broad substrate specificity [70]. Given that different AHLs are produced and sensed by a variety of bacteria [66,71–74], the use of these enzymes may capture signal-specific changes to the oral community. Lactonases reduce the pathogenicity of Gram-negative bacterial species both *in vitro* [75,76] and *in vivo* [77–80] and their substrate preference correlates with distinct proteome profiles, virulence factor expression, antibiotic resistance profiles, and biofilm formation [16,81,82].

Here, we investigated the presence of AHLs in different cultures of model human dental plaque communities [83,84] using plasmid-based biosensors. Remarkably, AHLs are detected in aerobic but not in anaerobic cultures. We then evaluated the effects of addition of exogenous AHLs of varying acyl chain lengths (C6 and C12) and treatment with QQ lactonases with distinct substrate preference. In response to these treatments, the microbial community population structure changed as determined using 16S rRNA sequencing. Interestingly, AHL signaling increases the relative abundance of bacterial taxa commonly considered as late colonizer pathogens associated with periodontal diseases, such as *Porphyromonas* and *Veillonella*. Conversely, the disruption of AHL signaling in dental plaque by QQ lactonases increased the abundance of bacterial taxa such as Lactobacillales, *Actinomyces* and *Streptococcus* that are dominant in a healthy oral microbiota and are considered to be pioneer colonizers of the tooth surface. Taken together, these results suggest a critical importance for AI-1/ AHL-based QS in the pathogenic profile of the oral microbiome.

## Materials and Methods

### Reagents

Chemicals including antibiotics were of reagent grade or superior and were sourced from Millipore Sigma (Burlington, MA) or Fisher Scientific (Hampton, NH). *N*-hexanoyl-L-homoserine lactone designated as C6-HSL (#10007896), *N*-3-oxo-octanoyl-L-Homoserine lactone designated as 3oC8-HSL (#10011206) and *N*-dodecanoyl-L-homoserine lactone designated as C12-HSL (#10011203) were purchased from Cayman Chemical Company (Ann Arbor, MI) and dissolved in 100% DMSO just before use.

### Lactonase production and purification

The *E. coli* strain BL21(DE3) was used to produce the inactive SsoPox mutant 5A8, hereafter referred to as 5A8, carrying the mutations V27G/P67Q/L72C/Y97S/Y99A/T177D/R223L/L226Q/L228M/W263H and was obtained previously [85], and SsoPox-W263I [69], hereafter referred to as SsoPox and GcL, respectively [70]. Purification was performed as previously described [70,82]. All lactonase preparations used in this study were made in buffer (PTE) composed of 50 mM HEPES pH 8.0, 150 mM NaCl, 0.2 mM CoCl_2_, sterilized by passing through a 0.2 μ filter, and stored at 4°C until use.

### *In vitro* dental plaque biofilms

#### Dental plaque community

We used a previously described representative supragingival plaque community (dental plaque) [83,84] as a model system to assess production of AHLs and their role in community composition. For the experiments described here, frozen 10% glycerol stock of dental plaque was used to inoculate fresh modified Shi medium (25% Shi medium and 75% of sterile human saliva) and incubated overnight at 37°C under 5% CO_2_ (we sometimes referred this culture condition in 5% CO_2_ as “aerobic” for simplification purposes). Cultures in CO_2_ were incubated in the presence of 200 μg/mL lactonases 5A8 (inactive lactonase; control), SsoPox or GcL. Anaerobic (10% H_2_, 10% CO_2_ and 80% N_2_) cultures included DMSO (1:1000 dilution; control), 10 μM C6-HSL or 10 μM C12-HSL as previously described [84]. Biofilms were grown overnight using saliva-coated-6-well polystyrene culture plates (Corning, NY).

#### Saliva collection and sterilization

Collection and sterilization of saliva was done as previously described [86]. Briefly, stimulated whole saliva was collected and pooled from at least three healthy volunteers using protocols that were reviewed and approved by the Institutional Review Boards from the University of Minnesota (STUDY00016289). Saliva was collected by expectoration into tubes on ice followed by centrifugation for 10 minutes at 4°C at 4000 *x g* (Beckman SX4750 rotor) to remove large debris and bacterial aggregates. The supernatants were collected, and the planktonic bacteria were pelleted by centrifugation at 15,000 *x g* for 20 minutes at 4°C. The supernatants were collected, and filter sterilized. Sterile saliva was mixed with Shi medium at 75:25 proportions of saliva to Shi medium.

### Detections of AHLs

The biosensor plasmid pJBA132 [87] contains a gfp(ASV) reporter gene under the transcriptional regulation of an AHL-responsive *luxI* promoter. pJBA132 was transformed into *E. coli* strain JM109 (Promega; Madison, WI). JM109 harboring pJBA132 was routinely cultured in Miller’s Luria-Bertani (LB) broth (BD Difco, #244610) or LB-agar plates (BD Difco, #244510, supplemented with 10 μg/mL tetracycline. An overnight culture of the biosensor strain derived from a single colony was diluted 1:100 into fresh LB broth supplemented with 10 μg/mL tetracycline and grown in a shaker incubator at 37°C and 225 rpm. At 2 hours, cultures reach an early exponential phase of growth. Cell-free culture supernatants of the dental plaque community were mixed with biosensor cultures at a ratio of either 1:4 or 1:5 and dispensed at a total volume of 200 μL per well (6 replicates) in specialized 96-well microplates designed for measuring fluorescence and optical density simultaneously (Grenier Bio-One^TM^ #655906, non-binding opaque black 96-well microplate with μClear™ Film Bottom). For the blank control, sterile, uninoculated modified Shi media was mixed with biosensor cultures at a similar dilution. The microplates were covered with a lid and incubated in a microplate shaker incubator (Titramax 1000 from heidolph instruments) at 37°C and 300 rpm for 3 hours. GFP fluorescence (Excitation: 485 nm. Emission: 528 nm; Gain 75) and OD_600nm_ were measured in a BioTek Synergy HTX microplate spectrophotometer. The data were represented as the ratio of arbitrary relative fluorescence units (RFUs) to the OD_600nm_ of the biosensor culture [RFU/OD_600nm_]. The “blank” was regarded as the mean RFU/OD_600nm_ of 6 replicates of “media-only” control and subtracted from the RFU/OD_600mn_ of all other results. After the subtraction of blank values, the RFU/OD_600nm_ was adjusted for dilution. To estimate the AHL concentrations detected by the biosensor, we used a standard curve determined with 3oC8-HSL (**Fig. S1**). For the biosensor standard curve, varying concentrations (1 nM – 1 mM in 1 log_10_ fold increments) of 3oC8-HSL freshly prepared in DMSO (2 µL) was diluted 1:100 to excite the exponentially growing biosensor strain culture (198 µL). Data was recorded as before and represented as the mean and error of RFU/OD_600nm_ of 6 biological replicates to generate the standard curve.

### DNA extraction and 16S sequencing

Total microbial DNA from biofilms and planktonic cells of dental plaque cultures was extracted using DNeasy PowerSoil kit (Qiagen; Germantown, MD). After the recommended quality control procedures, the DNA was submitted for sequencing of the V4 hypervariable region of 16S rRNA genes at the University of Minnesota’s Genomics Center (UMGC). 16S sequencing data were deposited to the Sequence Read Archive (SRA) under the accession number PRJNA1083492.

### Bacterial Community Processing

16S rRNA amplicon sequences were processed with Mothur v.1.41.1 [88]. Forward and reverse reads were merged into contigs and filtered with defined criteria as previously described [89]. Briefly, any sequences that were removed were based upon ambiguous bases longer than 300 bp and homopolymers ≥ 8 nts. The processed sequences were then aligned against the SILVA database v.132.4 [87], subject to a 2% pre-cluster [90]. UCHIME was used to remove chimera within sequences [91]. Operational taxonomic units (OTUs) were classified at 97% sequence similarity using the furthest-neighbor algorithm and against the Ribosomal Database Project v.18 [92]. The collapsed taxon table was converted to relative abundance using the phyloseq package in RStudio. Alpha diversity metrics (Good’s coverage, Sob, Shannon, Chao, ACE, and Simpson) were calculated using the normalized OTU table, and Bray-Curtis similarities matrices were used for beta diversity comparisons in Mothur v.1.41.1 [88].

### Graphing and Statistical Analysis

Beta diversity comparisons, principal-coordinate and Lefse analyses were performed using Mothur v1.41.1 [88]. Differences in beta diversity between samples were analyzed using analysis of similarity (ANOSIM) [93] and analysis of molecular variances (ANOVA) [94]. Student’s t-test [95] was performed in R to calculate the differences in alpha diversity metrics, and Wilcoxon test [95] was used for the differences in OTU-level abundances between samples. Ggplot2 package in R was used to visualize the taxa abundances and diversities in bacterial communities between the samples [96]. Krona pie chart tool in Galaxy (https://github.com/marbl/Krona/wiki) was also used for visualization of Mothur results. The results from the biosensor experiments were processed, analyzed, and graphed using Microsoft Excel and GraphPad Prism 8. Unpaired two-tailed t-tests with Welch’s correction were used for determining statistical significance and calculated using GraphPad Prism. All the raw experimental data and analysis for this study is available at the Open Science Foundation at this <u>link</u>.

## Results

### AHLs are present in dental plaque cultures in CO_2_ atmosphere but not in anaerobic conditions

AHLs were previously reported to be produced by co-cultures of *Porphyromonas gingivalis* with *Streptococci* [53], two bacteria commonly associated with human dental plaque. Thus, we hypothesized that dental plaque communities could produce AHLs. We used microbial engineered biosensors to evaluate the AHL content of a previously characterized complex human oral microbial community derived from dental plaque [83,84]. *E. coli* strain JM109 harboring biosensor plasmid pJBA132 [87] was used to reliably quantify AHLs in the range of 10 - 100 nM, as shown by a standard titration with 3oC8-HSL (**Fig. S1**). Dental plaque community cultures grown under 5% CO_2_ produced ∼12.5 nM of AHLs (**Fig. 1 and Fig. S2),** a level consistent with previous reports. For example, co-cultures of *P. gingivalis* and *S. oralis* produce an estimated ∼5.8 nM of 3oC8-HSL [53]. To reduce the concentration of the produced AHLs in the cultures, lactonases SsoPox and GcL were used to corroborate the estimates. AHL concentrations are significantly decreased by 65% upon treatment with SsoPox (∼4.38 nM of AHLs) and 36% with GcL (∼7.96 nM of AHLs), respectively (**Fig. 1**). Given the substrate preference of SsoPox, long chain AHLs (C8 and above) appear to be produced in the tested cultures. Remarkably, AHLs were not detected in anaerobic cultures (**Fig. S3**).

**Fig. 1.**
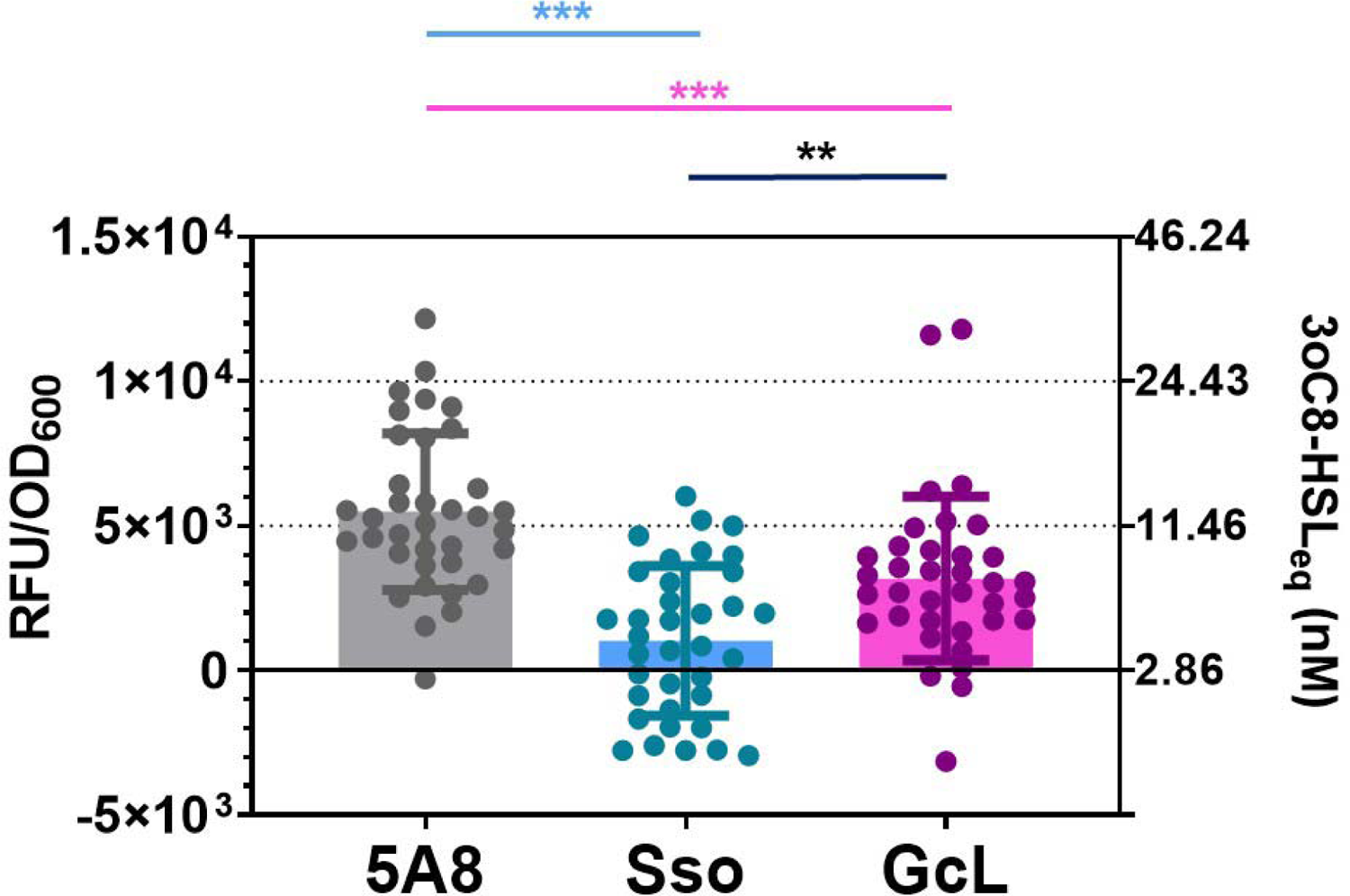
AHLs are detected in the spent supernatants of a dental plaque community cultured under 5% CO_2_ atmosphere or anaerobically. Cell-free culture supernatants of 6 biological replicates of aerobically cultured dental plaque community treated with lactonases 5A8 (inactive lactonase mutant), SsoPox, or GcL were incubated with *E. coli* AHL biosensor strain JM109 pJBA132. The resulting fluorescence is indicated as relative fluorescence units (RFU) per unit OD_600nm_ of biosensor cultures on the left Y-axis. The equivalent 3oC8-HSL concentration for the corresponding RFU/OD_600_ values as shown on the right Y-axis was interpolated from the standard curve of the biosensor response against 3oC8-HSL in **Fig. S1**. Results represent the mean and standard deviation of combined 36 replicates for each lactonase treatment, as each of the cell-free supernatants of 6 biological replicate cultures of a same model dental plaque community were further incubated with 6 biological replicates of biosensor cultures. Statistical significance of all treatments compared to the control (5A8) was calculated using unpaired two-tailed t-tests with Welch’s correction and significance values are indicated as - \*\*\**p < 0.0005*, \*\**p < 0.005* and \**p < 0.05*.

### CO_2_ atmosphere conditions affect the population structure of model dental plaque

Supragingival plaque microbial population structure has been previously well documented [97–99]. However, its potential changes induced by oxygen conditions are unclear. This is important because the oxygen levels vary greatly within the dental plaque structure and decrease sharply with plaque thickness [100]. We analyzed microbial population structures of supragingival plaque community grown in either 5% CO_2_ or anaerobic conditions. The CO_2_ and anaerobic communities show very different beta diversity (**Fig. 2A**). Furthermore, the inter-group differences between 5% CO_2_ and anaerobic communities were significantly greater than the intra-group differences (ANOSIM: R = 1, *p* = 0.003 (biofilm); R =1, *p* = 0.006 (planktonic)). To explore the taxa that best discriminate the 5% CO_2_ and anaerobic populations, we applied the linear discriminant analysis (LDA) effect size (LEfSe) method [101] to the biofilm samples (**Fig. 2B**). An LDA score was assigned to each taxonomic feature analyzed for differential abundance across different groups. Positive LDA scores were found for genera that were more prevalent in dental plaque samples grown under anaerobic conditions. Samples grown under 5% CO_2_ showed a higher prevalence of negative LDA scores. Four CO_2_ and nine anaerobic nested taxa best discriminated the two culture conditions. In the 5% CO_2_ condition, the key taxa are mainly Gram-positive bacteria such as *Abiotrophia*, *Schaalia* (formerly known as *Actinomyces*), Lactobacillales, and *Streptococcus*. Conversely, the key differentiating taxa in the anaerobic community were Gram-negative bacteria, including *Fusobacterium*, *Prevotella*, *Porphyromonas*, *Haemophilus*, and *Veillonella.* In general, we noticed that in all the treatments, Gram-positive taxa, including *Streptococcus*, *Peptostreptococcus*, *Parvimonas* and *Gemella* (32% in read abundance), were less abundant in both anaerobic biofilm and planktonic plaque communities than Gram-negative taxa *Fusobacterium*, *Prevotella*, *Porphyromonas*, *Haemophilus*, and *Veillonella* (42% in read abundance) (**Fig. S4A**). In contrast, CO_2_ samples showed Gram-negative taxa (*Fusobacterium*, *Prevotella*, and *Veilonella*) to represent less than 5% of the communities (**Figure S4B**). Planktonic cells from the plaque cultures behaved similarly in 5% CO_2_ and anaerobic conditions (**Fig. S5**). The anaerobic microbial population does not reveal many species that are reported to produce AHLs, except for *P. gingivalis* that can make small amounts of AHLs [52]. Most identified microbes were reported to produce AI-2 and/or AIPs [36,46,47,49,105–107]. Therefore, there may be production of AHLs by unknown sources or minor taxa within these communities.

**Fig. 2.**
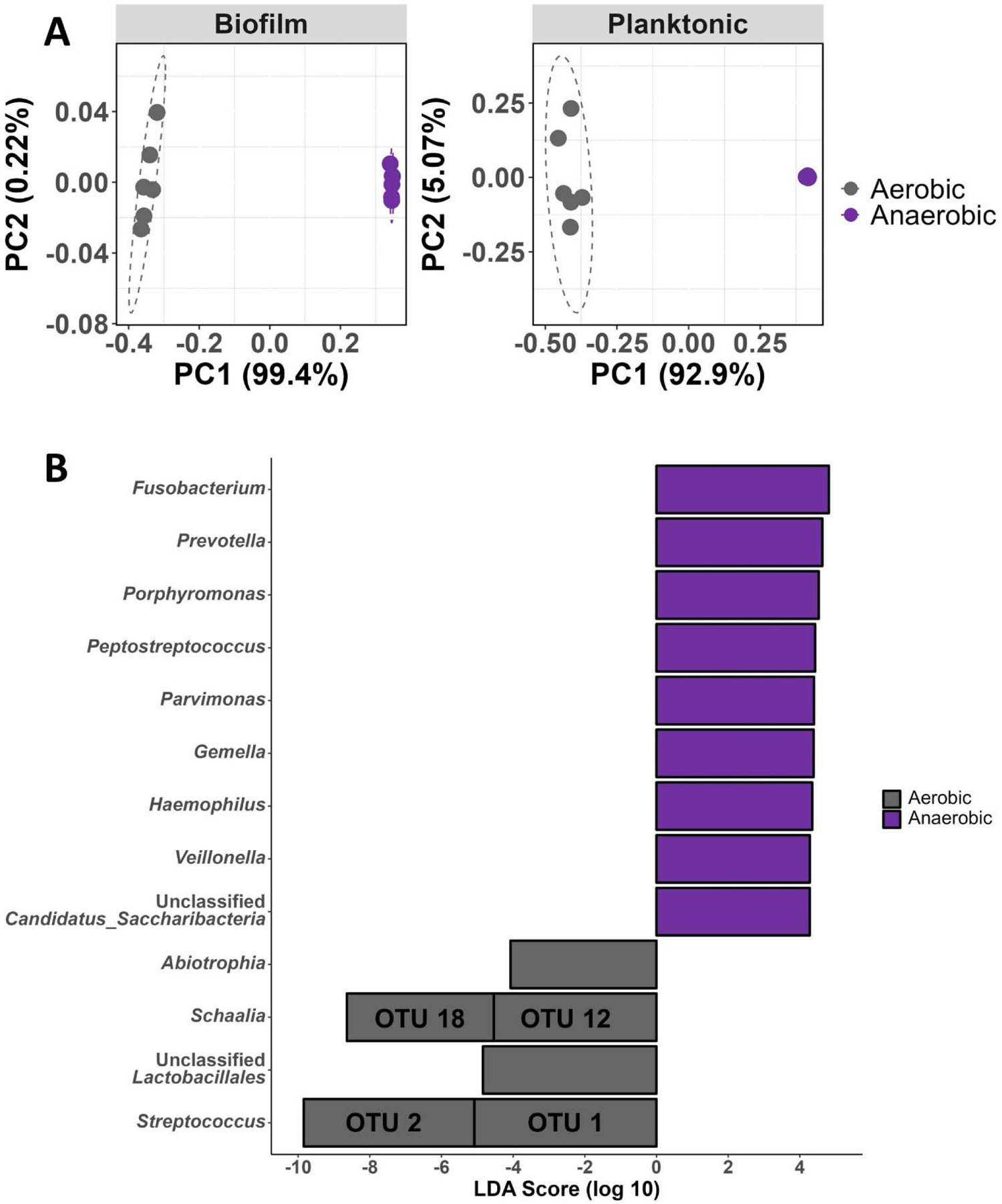
Microbial composition of a model dental plaque community under 5% CO_2_ and anaerobic culture conditions. **(A)** Principal-coordinate analysis (PCoA) of Bray-Curtis distances of both biofilm and planktonic dental plaque communities. **(B)** Linear discriminant analysis effect size (LEfSe) of biofilm dental plaque communities. The bar graph of LDA scores shows the taxa (OTU: Operational Taxonomic Unit) with statistical difference between 5% CO_2_ and anaerobic communities. Only taxa meeting a LDA significant threshold > 4 are shown.

### AHL signal disruption by lactonases increased the abundance of commensals and early colonizers in dental plaque biofilms

The importance of AHL signaling to the microbial population structure was examined using QQ lactonases to disrupt AHL signaling and followed by 16S rRNA sequencing. Under 5% CO_2_, treatment with each lactonase yielded similarly abundant taxa in the biofilm and planktonic communities: *Streptococcus* (∼49.6% - 54.7%), followed by Lactobacillales (order) (∼17% - 21.6%), *Actinomyces* (∼9.6% - 18.9%), and *Abiotrophia* (∼8.3% - 11.6%) (**Fig. 3A and S6A**). These Gram-positive taxa are mostly commensals and early colonizers of the oral biofilms [102,103]. We also observed a large number of less abundant, phylogenetically distinct taxa in CO_2_ and anaerobic growth conditions (**Fig. S4**). These changes to the planktonic communities upon treatments with lactonases are moderately statistically significant when individually compared to control (**Table S1).** For example, treatment with GcL results in a community that is not robustly statistically different from control (5A8; AMOVA: *p* = 0.371; ANOSIM: R = 255; *p* = 0.007*; **Table S1 and S2**). Although community population changes are small, when analyzed together, population changes induced by lactonases are significant (AMOVA: *p* = 0.047*; ANOSIM: R = 0.305; *p* = 0.003*; **Table S1 and S2**). In addition, the alpha diversity was significantly decreased with the SsoPox lactonase treatment when compared to control (t-test: *p* = 0.013) and GcL lactonase (t-test: *p* = 0.031) (**Table S3**, **Fig. 3B and 3C**).

**Fig. 3.**
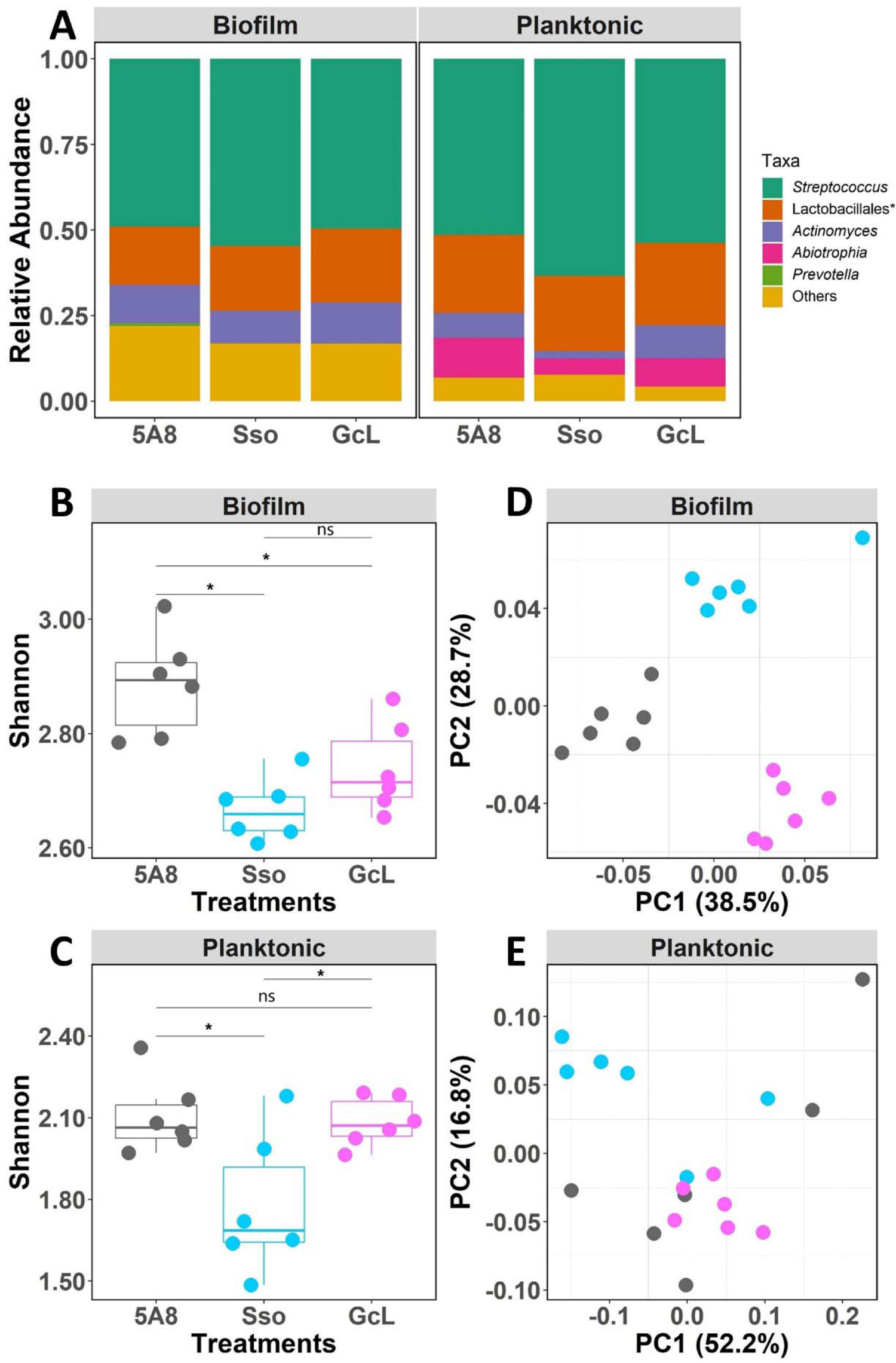
Lactonases altered the biofilm and planktonic oral microbiome of a dental plaque community cultured under 5% CO_2_ atmosphere conditions *in vitro*. (A) Relative abundance of the bacteria taxa at the genus level. “Others” represent taxa comprising less than 5% of the total relative abundance per sample. **(B)** Alpha diversity (Shannon index) of the biofilm community colored by treatment (gray: SsoPox 5A8 (control); blue: SsoPox; pink: GcL). **(C)** Principal-coordinate analysis (PCoA) of Bray-Curtis distances of the biofilm dental plaque community. (**D**) Alpha diversity (Shannon index) of the planktonic community colored by treatment (gray: SsoPox 5A8 (control; blue: SsoPox; pink: GcL). Differences in the alpha diversity metrics by treatments were tested for by using a Student’s t-test, and significance values are indicated as \*\*\**p < 0.0005*, \*\**p < 0.005* and \**p < 0.05, ns p > 0.05.* (E) Principal-coordinate analysis (PCoA) of Bray-Curtis distances of the planktonic dental plaque community.

On the other hand, the disruption of AHL signaling has much larger effects on the sessile microbial population **(Table S1 and S2**). SsoPox and GcL lactonase treatments significantly altered the microbial community composition (AMOVA: *p* = 0.002*, ANOSIM: R = 1, *p* =0.002* (control vs GcL); AMOVA: *p* = 0.002*, ANOSIM: R = 0.733, *p* = 0.002* (control vs SsoPox)). At the genus level, treatment with GcL resulted in significantly increased relative abundance averages of Lactobacillales (∼21.6%) and *Actinomyces* (∼12.1%), compared to control (∼17.0% and ∼11.2%, respectively; Wilcox test, *p* < 0.01). Treatment with the lactonase SsoPox also resulted in community changes, with a statistically significant increase of Lactobacillales (∼18.9%) and *Streptococcus* (∼54.7%), but a decrease in the relative abundance of *Actinomyces* (∼9.6%; Wilcox test, *p* < 0.01, **Fig. S7A**). SsoPox and GcL lactonase treatments increased the total relative abundance of these commensal and early colonizer Gram-positive bacterial taxa (Lactobacillales, *Actinomyces, and Streptococcus*) by ∼6% (to ∼83%; from ∼77.2% in control). The alpha diversity (Shannon index) was significantly lower with either GcL or SsoPox treatments compared to control (t-test: *p* = 0.013, *p* = 0.00083, respectively; **Table S3**, **Fig. 3B and 3C**). The community composition as a whole (beta diversity) exhibited differences between the two lactonase treatments and between the lactonase treatments and control group in both biofilm and planktonic communities. These differences were reflected in distinct clustering patterns on the principal-coordinate analysis (PCoA) of Bray-Curtis distances (**Fig. 3D and 3E**). Thus, AHL signal disruption using lactonases appears to alter the diversity and composition of both the biofilm (sessile) and planktonic communities under 5% CO_2_ atmosphere conditions. These changes are much larger in the sessile population, highlighting the importance of AHL signaling in these communities.

Interestingly, the treatments with the two distinct lactonases also lead to different communities in both culture conditions (AMOVA: p < 0.001*, ANOSIM: R = 0.83, p < 0.001* (biofilm); AMOVA: p = 0.018, ANOSIM: R = 0.51, p = 0.01* (planktonic)) (**Table S1 and S2**) and different diversity of the major bacterial genera. Unexpectedly, some of the taxa that were most affected by lactonases (e.g. Lactobacillales, *Actinomyces*, and *Streptococcus*) are Gram-positive bacteria and are not known to produce or use AHLs. Specifically, in the context of biofilm communities, GcL treatment leads to a notable increase in the average relative abundance of *Actinomyces* (12.1%) and Lactobacillales (21.6%) compared to SsoPox treatment (9.6%, 18.9%, respectively). Conversely, only *Streptococcus* demonstrates a decreased average relative abundance in the presence of GcL (49.6%) compared to SsoPox (54.7%) (**Figure S7A**). However, there were no significant differences between the treatments in the planktonic communities (**Figure S7B**). These differences in the plaque microbial community between the two lactonase treatments likely reflect the AHL substrate preferences of the two enzymes [69,70]. GcL is a broad spectrum AHL degrading enzyme [70] while SsoPox prefers long chains [69].

AHL disruption may have the potential to affect most bacteria within a community, whether or not they directly produce, use, or sense AHLs. If the effect is indirect, AHL disruption may alter the abundance and/or the metabolism of AHL-sensing bacteria, and these changes in turn influence other bacteria.

### Exogenous AHLs increased the abundance of late colonizers in anaerobic cultures of dental plaque biofilms

In contrast to the cultures grown under 5% CO_2_, no AHLs were detected under anaerobic conditions. Building on this observation, we assessed how the addition of exogenous AHLs (*i.e.* C6-HSL or C12-HSL) could affect community composition, particularly the major taxa in both planktonic and sessile samples (**Fig. 4A and S6B**).

**Fig. 4.**
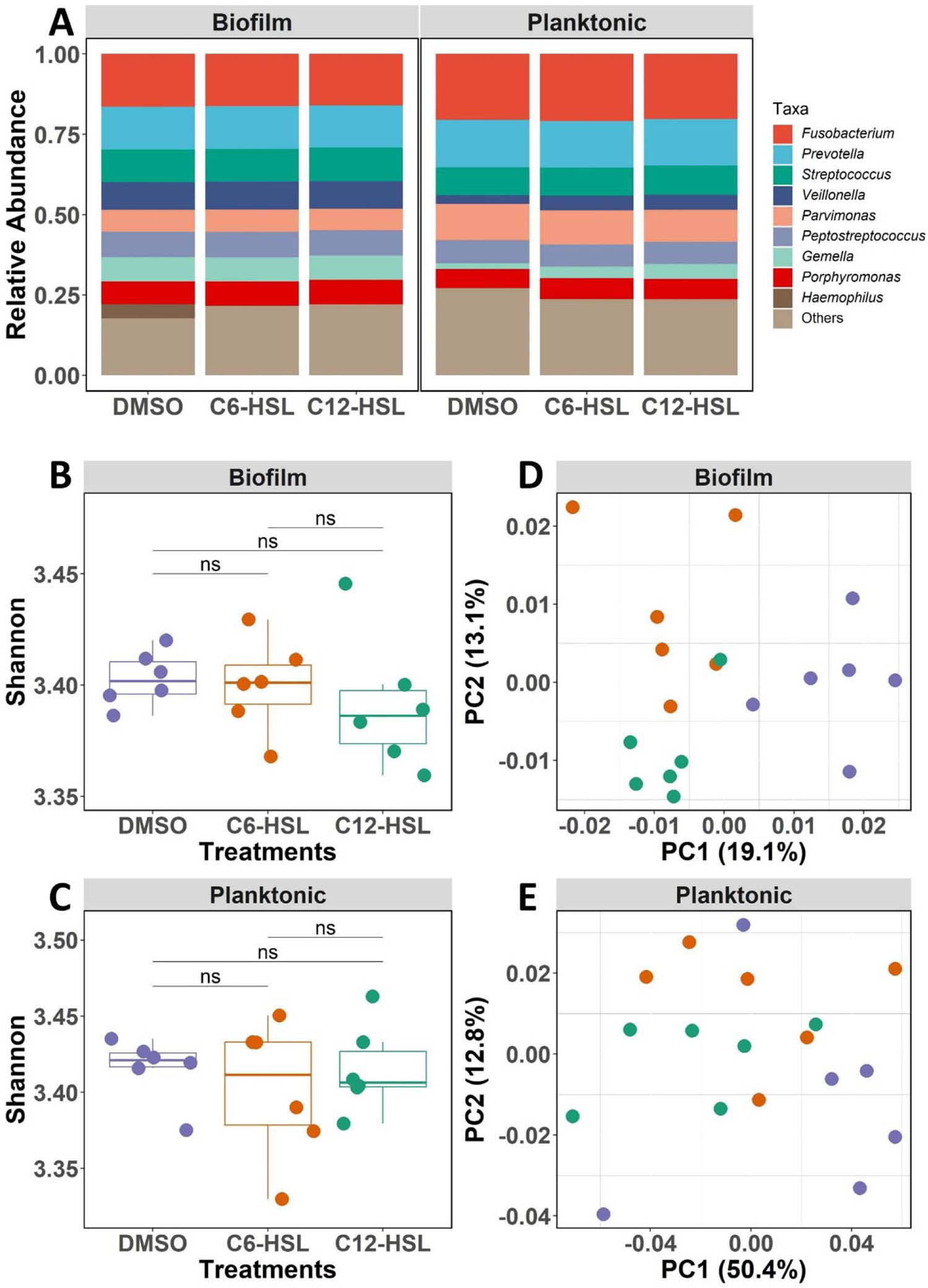
AHLs altered the biofilm and planktonic oral microbiomes of a dental plaque community cultured under anaerobic conditions *in vitro*. (A) Relative abundance of the bacteria taxa at the genus level. “Others” represent taxa comprising less than 5% of the total relative abundance per sample. **(B)** Alpha diversity (Shannon index) colored by treatment (grey: DMSO (control); blue: C6-HSL, pink: C12-HSL). Differences in the alpha diversity metrics by treatments were tested for by using a Student’s t-test, and significance values are indicated as - \*\*\**p < 0.0005*, \*\**p < 0.005* and \**p < 0.05, ns p > 0.05*. **(C)** Principal-coordinate analysis (PCoA) of Bray-Curtis distances of the biofilm dental plaque communities. (**D**) Alpha diversity (Shannon index) of the planktonic community colored by treatment (gray: SsoPox 5A8 (control; blue: SsoPox; pink: GcL). Differences in the alpha diversity metrics by treatments were tested for by using a Student’s t-test, and significance values are indicated as - \*\*\**p < 0.0005*, \*\**p < 0.005* and \**p < 0.05, ns p > 0.05.* (E) Principal-coordinate analysis (PCoA) of Bray-Curtis distances of the planktonic dental plaque community.

AHL treatments did not significantly alter the alpha diversity of sessile biofilm or planktonic samples (**Fig. 4B and 4C**). However, addition of AHLs significantly alters the sessile biofilm communities (AMOVA: *p* < 0.001*; ANOSIM: R = 0.54, *p* < 0.001*), but not the planktonic populations (AMOVA: *p* = 0.126; ANOSIM: *p* = 0.071) (**Fig. 4D and 4E; Table S4 and S5**). Interestingly, *Haemophilus* (∼5.3%), a commensal Gram-negative bacterium that can have positive immunomodulatory effects [104,105], was detected only in control biofilm samples, but not in AHL-treated samples (**Fig. 4A and S6B**). The examination of the changes caused by the addition of C12-HSL reveal that anaerobic sessile biofilms showed a decrease in the relative abundance of *Fusobacterium* (∼16.0%, as compared to ∼16.5% in control) and increased abundance of *Porphyromonas* (∼7.7%; ∼7.0% in control) (Wilcox test: *p* = 0.0043) (**Fig. S7C**). The treatment with C6-HSL resulted in a significant increase of *Porphyromonas* (∼7.6%; ∼7.0% in control) and *Veillonella* (∼8.8%; 8.5% in control) (Wilcox test: *p* = 0.015, **Fig. S7C**). Addition of AHLs had been reported to increase periodontal pathogens such as *Fusobacterium* and *Porphyromonas* [51]. In anaerobic samples, the planktonic community showed an increase in the relative abundance of *Streptococcus* when C12-HSL was added (9.1%) that was significantly higher compared to C6-HSL (∼8.7%) and control treatments (∼8.6%) (Wilcox test: *p* = 0.026, **Fig. S7D**). No other significant differences in bacteria taxa were found between control and AHL-treated anaerobic planktonic cultures. Therefore, signal induction using exogenous AHLs appears to alter the diversity and composition of both the biofilm and sessile communities grown anaerobically, with distinct outcomes depending on their acyl group chemistry, particularly increasing the abundance of late colonizer species and periodontal pathogens in the plaque biofilms.

## Discussion

The microbial population structure of supragingival dental plaque has been well characterized [97–99,102,103,106]. The pioneer and early colonizers are primarily composed of Streptococci, Lactobacillales and *Actinomyces,* whose abundance is in indicator of a healthy oral microbiota. Subsequent colonizers include *Capnocytophaga*, *Eiknella, Veillonella, Fusobacterium*, *Prevotella*, *Campylobacter, Porphyromonas gingivalis*, *Tannerella forsythia* and *Treponema Denticola*, some of which are well documented periodontal pathogens. These Gram-positive and Gram-negative taxa are not known to produce AHLs, except for *P. gingivalis* that can make small amounts of AHLs [52] and were reported to produce AI-2 and/or AIPs [36,46,47,49,105–107].

The role(s) of AHL signaling in oral communities have remained elusive. The presence of AHLs itself is heavily discussed [53]. In fact, AHLs were previously undetected using *Escherichia coli, Chromobacterium violaceum* and *Vibrio harveyi* based biosensors in anaerobic axenic cultures of Gram-negative pathogens such as *Porphyromonas gingivalis*, *Veillonella parvula*, *Prevotella intermedia* and *Fusobacterium nucleatum* [37,47]. Here we find that AHLs could not be detected in anaerobic conditions but were abundant in dental plaque cultures in 5% CO_2_. This finding is consistent with observations made on other types of biofilms: the concentration gradient of AHL is in fact reported to coincide with the oxygen gradient [107]. This new finding is likely essential to understand the potential role(s) of AHL signaling in oral communities. As oxygen levels vary during biofilm growth and within sessile communities [108–110], the role of AHLs is likely to vary greatly. The role of AHLs could also vary as a reflection of the spatial location within the plaque.

This potential role(s) of AHL signaling in oral communities is further evidenced by the different effects of both AHL exogenous addition and signal disruption on plaque cultures grown in anaerobic and more aerobic conditions. For instance, in conditions where AHLs are detected (5% CO_2_), we observed that signal disruption using QQ lactonases results in large, significant changes in microbial populations of a dental plaque biofilm community. Despite the non-detectable level of AHLs in anaerobic conditions, we find that the addition of AHLs results in large changes in anaerobic biofilms. This is strong evidence that AHL signaling plays a significant role in maintaining the plaque community in both anaerobic and more aerobic conditions.

Interestingly, this work reveals that AHL signaling affects critical microbes for plaque establishment and oral health. For instance, AHL signal disruption in 5% CO_2_ cultures favors *Streptococcus,* a finding that resonates with previous reports [111,112]. Lactonase treatments also resulted in an increased abundance of commensals and early colonizers (e.g. Lactobacillales, *Streptococcus, Actinomyces)* at the expense of Gram-negative pathogens. Importantly, we find that changes induced to the community population structure depends on the AHL preference of the lactonase. This new finding simulates earlier reports in which lactonase was reported to reduce cariogenic and periodontal biofilms and the abundance of periodontal pathogens including *Porphyromonas gingivalis*, *Tannerella forsythia* and *Treponema Denticola* under anaerobic conditions [53,62,106,113]. Hence, AHL disruption favors growth of bacterial taxa associated with healthy oral microbiota.

Diverse AHL signals have been detected in oral biofilms [53,62,113], and here we highlight the importance of these signaling molecules by exogenously adding them to cultures. In anaerobic dental plaque cultures, the addition of both C6-HSL and C12-HSL leads to an increase in the abundance of *Porphyromonas*. When adding C6-HSL treatment, the abundance of *Veillonella* increases, whereas C12-HSL treatment decreased abundance of *Fusobacterium*. Overall, the presence of AHLs favors emergence of late colonizing periodontal pathogens in the sessile dental plaque community. Consistently with the observations from lactonase treatments, the changes induced to the community population structure depends on the chemical structure of the added AHLs.

Anaerobic communities fail to produce AHLs yet are responsive to these signals. Hence, the emergence of a periodontal pathogenic community could depend on the specific QS signaling molecules produced in more aerobic spatial locations in dental plaque. It is also possible that produce AHLs could diffuse from aerobic compartments and affect anaerobic colonizers. We successfully showed the role of QS in the behavior of a complex community including the selection of partners in the community. Such behaviors may not be simulated by simpler two-species or small microcosm systems. Altering microbial interactions to interfere in biofilm formation has been proposed as a prophylactic strategy for oral microbial infections [114]. Further studies will be needed to decipher the roles of these specific AHL signals in the different stages of oral biofilm and disease development.

## ACKNOWLEDGEMENTS

We would like to thank Dr. David Daudé at Gene&GreenTK for kindly providing the biosensor plasmid pJBA132. This work was conducted with support from the award no. R35GM133487 (to MHE) from the National Institute of General Medical Sciences and R01DE025618 from the National Institute of Dental and Craniofacial Research (MCH). MB was supported by a Biotechnology Training Grant: NIH T32GM008347 and UMII MnDRIVE Graduate Assistantship. The content is solely the responsibility of the authors and does not necessarily represent the official views of the National Institutes of Health.

## References

1. Miller MB, Bassler BL. Quorum sensing in bacteria. Annu Rev Microbiol. 2001;55:165–99.

2. Williams P. Quorum sensing, communication and cross-kingdom signalling in the bacterial world. Microbiology (Reading). 2007 Dec;153(Pt 12):3923–3938.

3. Papenfort K, Bassler BL. Quorum sensing signal-response systems in Gram-negative bacteria. Nat Rev Microbiol. 2016 Aug 11;14(9):576–88.

4. Biswa P, Doble M. Production of acylated homoserine lactone by gram-positive bacteria isolated from marine water. FEMS Microbiol Lett. 2013 Jun;343(1):34–41.

5. Zhang G, Zhang F, Ding G, et al. Acyl homoserine lactone-based quorum sensing in a methanogenic archaeon. ISME J. 2012 Jul;6(7):1336–44.

6. Pereira CS, Thompson JA, Xavier KB. AI-2-mediated signalling in bacteria. FEMS Microbiol Rev. 2013 Mar;37(2):156–81.

7. Nichols JD, Johnson MR, Chou CJ, et al. Temperature, not LuxS, mediates AI-2 formation in hydrothermal habitats. FEMS Microbiol Ecol. 2009 May;68(2):173–81.

8. Surette MG, Miller MB, Bassler BL. Quorum sensing in Escherichia coli, Salmonella typhimurium, and Vibrio harveyi: a new family of genes responsible for autoinducer production. Proc Natl Acad Sci U S A. 1999 Feb 16;96(4):1639–44.

9. Zhao L, Xue T, Shang F, et al. *Staphylococcus aureus* AI-2 quorum sensing associates with the KdpDE two-component system to regulate capsular polysaccharide synthesis and virulence. Infection and immunity. 2010 Aug;78(8):3506–15.

10. Chen X, Schauder S, Potier N, et al. Structural identification of a bacterial quorum-sensing signal containing boron. Nature. 2002 Jan 31;415(6871):545-9.

11. Monnet V, Juillard V, Gardan R. Peptide conversations in Gram-positive bacteria. Crit Rev Microbiol. 2016 May;42(3):339–51.

12. Le KY, Otto M. Quorum-sensing regulation in staphylococci-an overview. Front Microbiol. 2015;6:1174.

13. Slamti L, Lereclus D. Specificity and polymorphism of the PlcR-PapR quorum-sensing system in the Bacillus cereus group. J Bacteriol. 2005 Feb;187(3):1182–7.

14. Liu H, Coulthurst SJ, Pritchard L, et al. Quorum sensing coordinates brute force and stealth modes of infection in the plant pathogen *Pectobacterium atrosepticum*. PLoS pathogens. 2008 Jun 20;4(6):e1000093.

15. Mukherjee S, Bassler BL. Bacterial quorum sensing in complex and dynamically changing environments. Nat Rev Microbiol. 2019 Jun;17(6):371–382.

16. Sikdar R, Elias MH. Evidence for Complex Interplay between Quorum Sensing and Antibiotic Resistance in Pseudomonas aeruginosa. Microbiol Spectr. 2022 Dec 21;10(6):e0126922.

17. Montgomery K, Charlesworth JC, LeBard R, et al. Quorum sensing in extreme environments. Life (Basel). 2013 Jan 29;3(1):131–48.

18. Seneviratne CJ, Zhang CF, Samaranayake LP. Dental plaque biofilm in oral health and disease. Chin J Dent Res. 2011;14(2):87–94.

19. Human Microbiome Project C. Structure, function and diversity of the healthy human microbiome. Nature. 2012 Jun 13;486(7402):207–14.

20. Zaura E, Keijser BJ, Huse SM, et al. Defining the healthy “core microbiome” of oral microbial communities. BMC Microbiol. 2009 Dec 15;9:259.

21. Dewhirst FE, Chen T, Izard J, et al. The human oral microbiome. J Bacteriol. 2010 Oct;192(19):5002–17.

22. Willis JR, Gabaldon T. The Human Oral Microbiome in Health and Disease: From Sequences to Ecosystems. Microorganisms. 2020 Feb 23;8(2).

23. Perera M, Al-Hebshi NN, Speicher DJ, et al. Emerging role of bacteria in oral carcinogenesis: a review with special reference to perio-pathogenic bacteria. J Oral Microbiol. 2016;8:32762.

24. Zhao H, Chu M, Huang Z, et al. Variations in oral microbiota associated with oral cancer. Sci Rep. 2017 Sep 18;7(1):11773.

25. Belda-Ferre P, Alcaraz LD, Cabrera-Rubio R, et al. The oral metagenome in health and disease. ISME J. 2012 Jan;6(1):46–56.

26. Hajishengallis G, Darveau RP, Curtis MA. The keystone-pathogen hypothesis. Nat Rev Microbiol. 2012 Oct;10(10):717–25.

27. Marsh PD. Microbial ecology of dental plaque and its significance in health and disease. Adv Dent Res. 1994 Jul;8(2):263–71.

28. Mohanty R, Asopa SJ, Joseph MD, et al. Red complex: Polymicrobial conglomerate in oral flora: A review. J Family Med Prim Care. 2019 Nov;8(11):3480–3486.

29. Marsh PD. Dental plaque as a biofilm and a microbial community - implications for health and disease. BMC Oral Health. 2006 Jun 15;6 Suppl 1(Suppl 1):S14.

30. He X, Hu W, Kaplan CW, et al. Adherence to streptococci facilitates Fusobacterium nucleatum integration into an oral microbial community. Microb Ecol. 2012 Apr;63(3):532–42.

31. Kolenbrander PE, Palmer RJ, Jr., Rickard AH, et al. Bacterial interactions and successions during plaque development. Periodontol 2000. 2006;42:47–79.

32. Sakanaka A, Kuboniwa M, Shimma S, et al. Fusobacterium nucleatum Metabolically Integrates Commensals and Pathogens in Oral Biofilms. mSystems. 2022 Aug 30;7(4):e0017022.

33. Bramhachari PV, Ahmed VKS, Selvin J, et al. Quorum Sensing and Biofilm Formation by Oral Pathogenic Microbes in the Dental Plaques: Implication for Health and Disease. In: Bramhachari PV, editor. Implication of Quorum Sensing and Biofilm Formation in Medicine, Agriculture and Food Industry. Singapore: Springer Singapore; 2019. p. 129–140.

34. Wright PP, Ramachandra SS. Quorum Sensing and Quorum Quenching with a Focus on Cariogenic and Periodontopathic Oral Biofilms. Microorganisms. 2022 Sep 3;10(9).

35. Cuadra-Saenz G, Rao DL, Underwood AJ, et al. Autoinducer-2 influences interactions amongst pioneer colonizing streptococci in oral biofilms. Microbiology (Reading). 2012 Jul;158(Pt 7):1783–1795.

36. McNab R, Ford SK, El-Sabaeny A, et al. LuxS-based signaling in Streptococcus gordonii: autoinducer 2 controls carbohydrate metabolism and biofilm formation with Porphyromonas gingivalis. J Bacteriol. 2003 Jan;185(1):274–84.

37. Burgess NA, Kirke DF, Williams P, et al. LuxS-dependent quorum sensing in Porphyromonas gingivalis modulates protease and haemagglutinin activities but is not essential for virulence. Microbiology (Reading). 2002 Mar;148(Pt 3):763–772.

38. Rickard AH, Palmer RJ, Jr., Blehert DS, et al. Autoinducer 2: a concentration-dependent signal for mutualistic bacterial biofilm growth. Mol Microbiol. 2006 Jun;60(6):1446–56.

39. Fong KP, Chung WO, Lamont RJ, et al. Intra- and interspecies regulation of gene expression by Actinobacillus actinomycetemcomitans LuxS. Infection and immunity. 2001 Dec;69(12):7625–34.

40. Merritt J, Qi F, Goodman SD, et al. Mutation of luxS affects biofilm formation in Streptococcus mutans. Infection and immunity. 2003 Apr;71(4):1972–9.

41. Jang YJ, Choi YJ, Lee SH, et al. Autoinducer 2 of Fusobacterium nucleatum as a target molecule to inhibit biofilm formation of periodontopathogens. Arch Oral Biol. 2013 Jan;58(1):17–27.

42. Jang YJ, Sim J, Jun HK, et al. Differential effect of autoinducer 2 of Fusobacterium nucleatum on oral streptococci. Arch Oral Biol. 2013 Nov;58(11):1594–602.

43. Petersen FC, Pecharki D, Scheie AA. Biofilm mode of growth of Streptococcus intermedius favored by a competence-stimulating signaling peptide. J Bacteriol. 2004 Sep;186(18):6327–31.

44. Senadheera D, Cvitkovitch DG. Quorum sensing and biofilm formation by Streptococcus mutans. Advances in experimental medicine and biology. 2008;631:178–88.

45. Perry JA, Cvitkovitch DG, Levesque CM. Cell death in Streptococcus mutans biofilms: a link between CSP and extracellular DNA. FEMS Microbiol Lett. 2009 Oct;299(2):261–6.

46. Senadheera D, Krastel K, Mair R, et al. Inactivation of VicK affects acid production and acid survival of Streptococcus mutans. J Bacteriol. 2009 Oct;191(20):6415–24.

47. Frias J, Olle E, Alsina M. Periodontal pathogens produce quorum sensing signal molecules. Infection and immunity. 2001 May;69(5):3431–4.

48. Whittaker CJ, Klier CM, Kolenbrander PE. Mechanisms of adhesion by oral bacteria. Annu Rev Microbiol. 1996;50:513–52.

49. Guo L, He X, Shi W. Intercellular communications in multispecies oral microbial communities. Front Microbiol. 2014;5:328.

50. Huang R, Li M, Gregory RL. Bacterial interactions in dental biofilm. Virulence. 2011 Sep-Oct;2(5):435–44.

51. Muras A, Mayer C, Otero-Casal P, et al. Short-Chain N-Acylhomoserine Lactone Quorum-Sensing Molecules Promote Periodontal Pathogens in In Vitro Oral Biofilms. Applied and environmental microbiology. 2020 Jan 21;86(3).

52. Kumari A, Pasini P, Daunert S. Detection of bacterial quorum sensing N-acyl homoserine lactones in clinical samples. Anal Bioanal Chem. 2008 Jul;391(5):1619–27.

53. Muras A, Otero-Casal P, Blanc V, et al. Acyl homoserine lactone-mediated quorum sensing in the oral cavity: a paradigm revisited. Sci Rep. 2020 Jun 17;10(1):9800.

54. Chen JW, Chin S, Tee KK, et al. N-acyl homoserine lactone-producing Pseudomonas putida strain T2-2 from human tongue surface. Sensors (Basel). 2013 Sep 30;13(10):13192–203.

55. Yin WF, Purmal K, Chin S, et al. Long chain N-acyl homoserine lactone production by Enterobacter sp. isolated from human tongue surfaces. Sensors (Basel). 2012 Oct 24;12(11):14307–14.

56. Yin WF, Purmal K, Chin S, et al. N-acyl homoserine lactone production by Klebsiella pneumoniae isolated from human tongue surface. Sensors (Basel). 2012;12(3):3472–83.

57. Goh SY, Khan SA, Tee KK, et al. Quorum sensing activity of Citrobacter amalonaticus L8A, a bacterium isolated from dental plaque. Sci Rep. 2016 Feb 10;6:20702.

58. Goh SY, Tan WS, Khan SA, et al. Unusual multiple production of N-acylhomoserine lactones a by Burkholderia sp. strain C10B isolated from dentine caries. Sensors (Basel). 2014 May 21;14(5):8940–9.

59. Wu J, Lin X, Xie H. Regulation of hemin binding proteins by a novel transcriptional activator in Porphyromonas gingivalis. J Bacteriol. 2009 Jan;191(1):115–22.

60. Chawla A, Hirano T, Bainbridge BW, et al. Community signalling between Streptococcus gordonii and Porphyromonas gingivalis is controlled by the transcriptional regulator CdhR. Mol Microbiol. 2010 Dec;78(6):1510–22.

61. Aleti G, Baker JL, Tang X, et al. Identification of the Bacterial Biosynthetic Gene Clusters of the Oral Microbiome Illuminates the Unexplored Social Language of Bacteria during Health and Disease. mBio. 2019 Apr 16;10(2).

62. Parga A, Muras A, Otero-Casal P, et al. The quorum quenching enzyme Aii20J modifies in vitro periodontal biofilm formation. Front Cell Infect Microbiol. 2023;13:1118630.

63. Billot R, Plener L, Jacquet P, et al. Engineering acyl-homoserine lactone-interfering enzymes toward bacterial control. J Biol Chem. 2020 Sep 11;295(37):12993–13007.

64. Elias M, Tawfik DS. Divergence and convergence in enzyme evolution: parallel evolution of paraoxonases from quorum-quenching lactonases. J Biol Chem. 2012 Jan 2;287(1):11–20.

65. Remy B, Mion S, Plener L, et al. Interference in Bacterial Quorum Sensing: A Biopharmaceutical Perspective. Front Pharmacol. 2018;9:203.

66. Sikdar R, Elias M. Quorum quenching enzymes and their effects on virulence, biofilm, and microbiomes: a review of recent advances. Expert Rev Anti Infect Ther. 2020 Dec;18(12):1221–1233.

67. Elias M, Dupuy J, Merone L, et al. Structural basis for natural lactonase and promiscuous phosphotriesterase activities. J Mol Biol. 2008 Jun 20;379(5):1017–28.

68. Hiblot J, Gotthard G, Chabriere E, et al. Characterisation of the organophosphate hydrolase catalytic activity of SsoPox. Sci Rep. 2012;2:779.

69. Hiblot J, Gotthard G, Elias M, et al. Differential active site loop conformations mediate promiscuous activities in the lactonase SsoPox. PLoS One. 2013;8(9):e75272.

70. Bergonzi C, Schwab M, Naik T, et al. The Structural Determinants Accounting for the Broad Substrate Specificity of the Quorum Quenching Lactonase GcL. Chembiochem. 2019 Jul 15;20(14):1848–1855.

71. Liao L, Schaefer AL, Coutinho BG, et al. An aryl-homoserine lactone quorum-sensing signal produced by a dimorphic prosthecate bacterium. Proc Natl Acad Sci U S A. 2018 Jul 17;115(29):7587–7592.

72. Wellington S, Greenberg EP. Quorum Sensing Signal Selectivity and the Potential for Interspecies Cross Talk. mBio. 2019 Mar 5;10(2).

73. Brameyer S, Heermann R. Specificity of Signal-Binding via Non-AHL LuxR-Type Receptors. PLoS One. 2015;10(4):e0124093.

74. Grandclement C, Tannieres M, Morera S, et al. Quorum quenching: role in nature and applied developments. FEMS Microbiol Rev. 2016 Jan;40(1):86–116.

75. Guendouze A, Plener L, Bzdrenga J, et al. Effect of Quorum Quenching Lactonase in Clinical Isolates of *Pseudomonas aeruginosa* and Comparison with Quorum Sensing Inhibitors. Front Microbiol. 2017;8:227.

76. Lopez-Jacome LE, Garza-Ramos G, Hernandez-Duran M, et al. AiiM Lactonase Strongly Reduces Quorum Sensing Controlled Virulence Factors in Clinical Strains of *Pseudomonas aeruginosa* Isolated From Burned Patients. Front Microbiol. 2019;10:2657.

77. Bijtenhoorn P, Mayerhofer H, Muller-Dieckmann J, et al. A novel metagenomic short-chain dehydrogenase/reductase attenuates *Pseudomonas aeruginosa* biofilm formation and virulence on *Caenorhabditis elegans*. PLoS One. 2011;6(10):e26278.

78. Hraiech S, Hiblot J, Lafleur J, et al. Inhaled lactonase reduces *Pseudomonas aeruginosa* quorum sensing and mortality in rat pneumonia. PLoS One. 2014;9(10):e107125.

79. Stoltz DA, Ozer EA, Ng CJ, et al. Paraoxonase-2 deficiency enhances *Pseudomonas aeruginosa* quorum sensing in murine tracheal epithelia. Am J Physiol Lung Cell Mol Physiol. 2007 Apr;292(4):L852–60.

80. Utari PD, Setroikromo R, Melgert BN, et al. PvdQ Quorum Quenching Acylase Attenuates *Pseudomonas aeruginosa* Virulence in a Mouse Model of Pulmonary Infection. Front Cell Infect Microbiol. 2018;8:119.

81. Mahan K, Martinmaki R, Larus I, et al. Effects of Signal Disruption Depends on the Substrate Preference of the Lactonase. Front Microbiol. 2019;10:3003.

82. Remy B, Plener L, Decloquement P, et al. Lactonase Specificity Is Key to Quorum Quenching in *Pseudomonas aeruginosa*. Front Microbiol. 2020;11:762.

83. Hall JW, Lima BP, Herbomel GG, et al. An intramembrane sensory circuit monitors sortase A-mediated processing of streptococcal adhesins. Sci Signal. 2019 May 7;12(580).

84. Saavedra FM, Pelepenko LE, Boyle WS, et al. In vitro physicochemical characterization of five root canal sealers and their influence on an ex vivo oral multi-species biofilm community. Int Endod J. 2022 Jul;55(7):772–783.

85. Jacquet P, Hiblot J, Daude D, et al. Rational engineering of a native hyperthermostable lactonase into a broad spectrum phosphotriesterase. Sci Rep. 2017 Dec 1;7(1):16745.

86. Nairn BL, Lee GT, Chumber AK, et al. Uncovering Roles of Streptococcus gordonii SrtA-Processed Proteins in the Biofilm Lifestyle. J Bacteriol. 2020 Dec 18;203(2).

87. Andersen JB, Heydorn A, Hentzer M, et al. gfp-based N-acyl homoserine-lactone sensor systems for detection of bacterial communication. Applied and environmental microbiology. 2001 Feb;67(2):575–85.

88. Schloss PD, Westcott SL, Ryabin T, et al. Introducing mothur: open-source, platform-independent, community-supported software for describing and comparing microbial communities. Applied and environmental microbiology. 2009 Dec;75(23):7537–41.

89. Chappidi S, Villa EC, Cantarel BL. Using Mothur to Determine Bacterial Community Composition and Structure in 16S Ribosomal RNA Datasets. Curr Protoc Bioinformatics. 2019 Sep;67(1):e83.

90. Pruesse E, Quast C, Knittel K, et al. SILVA: a comprehensive online resource for quality checked and aligned ribosomal RNA sequence data compatible with ARB. Nucleic Acids Res. 2007;35(21):7188–96.

91. Edgar RC, Haas BJ, Clemente JC, et al. UCHIME improves sensitivity and speed of chimera detection. Bioinformatics. 2011 Aug 15;27(16):2194–200.

92. Cole JR, Wang Q, Cardenas E, et al. The Ribosomal Database Project: improved alignments and new tools for rRNA analysis. Nucleic Acids Res. 2009 Jan;37(Database issue):D141–5.

93. Clarke KR. Non-parametric multivariate analyses of changes in community structure. Australian Journal of Ecology. 1993;18(1):117–143.

94. Excoffier L, Smouse PE, Quattro JM. Analysis of molecular variance inferred from metric distances among DNA haplotypes: application to human mitochondrial DNA restriction data. Genetics. 1992 Jun;131(2):479–91.

95. Fay MP, Proschan MA. Wilcoxon-Mann-Whitney or t-test? On assumptions for hypothesis tests and multiple interpretations of decision rules. Stat Surv. 2010;4:1–39.

96. Hadley W. Ggplot2. New York, NY: Springer Science+Business Media, LLC; 2016.

97. de Jesus VC, Khan MW, Mittermuller BA, et al. Characterization of Supragingival Plaque and Oral Swab Microbiomes in Children With Severe Early Childhood Caries. Front Microbiol. 2021;12:683685.

98. Espinoza JL, Harkins DM, Torralba M, et al. Supragingival Plaque Microbiome Ecology and Functional Potential in the Context of Health and Disease. mBio. 2018 Nov 27;9(6).

99. Rabe A, Gesell Salazar M, Michalik S, et al. Impact of different oral treatments on the composition of the supragingival plaque microbiome. J Oral Microbiol. 2022;14(1):2138251.

100. Mangal U, Kwon JS, Choi SH. Bio-Interactive Zwitterionic Dental Biomaterials for Improving Biofilm Resistance: Characteristics and Applications. Int J Mol Sci. 2020 Nov 29;21(23).

101. Segata N, Izard J, Waldron L, et al. Metagenomic biomarker discovery and explanation. Genome Biol. 2011 Jun 24;12(6):R60.

102. Socransky SS, Haffajee AD, Cugini MA, et al. Microbial complexes in subgingival plaque. J Clin Periodontol. 1998 Feb;25(2):134–44.

103. Sukhvinder Singh O, Shabina S, Shibani G. Various complexes of the oral microbial flora in periodontal disease. Journal of Dental Problems and Solutions. 2021 2021/04/10:032-033.

104. Perera D, McLean A, Morillo-Lopez V, et al. Mechanisms underlying interactions between two abundant oral commensal bacteria. ISME J. 2022 Apr;16(4):948–957.

105. Tseng YC, Yang HY, Lin WT, et al. Salivary dysbiosis in Sjogren’s syndrome and a commensal-mediated immunomodulatory effect of salivary gland epithelial cells. NPJ Biofilms Microbiomes. 2021 Mar 11;7(1):21.

106. Carrouel F, Viennot S, Santamaria J, et al. Quantitative Molecular Detection of 19 Major Pathogens in the Interdental Biofilm of Periodontally Healthy Young Adults. Front Microbiol. 2016;7:840.

107. Flemming HC, Wingender J, Szewzyk U, et al. Biofilms: an emergent form of bacterial life. Nat Rev Microbiol. 2016 Aug 11;14(9):563–75.

108. Ahn SJ, Burne RA. Effects of oxygen on biofilm formation and the AtlA autolysin of Streptococcus mutans. J Bacteriol. 2007 Sep;189(17):6293–302.

109. Bradshaw DJ, Marsh PD, Allison C, et al. Effect of oxygen, inoculum composition and flow rate on development of mixed-culture oral biofilms. Microbiology (Reading). 1996 Mar;142 (Pt 3):623–629.

110. Mystkowska J, Niemirowicz-Laskowska K, Lysik D, et al. The Role of Oral Cavity Biofilm on Metallic Biomaterial Surface Destruction-Corrosion and Friction Aspects. Int J Mol Sci. 2018 Mar 6;19(3).

111. Tufekci EF, Akçay S, Kayıpmaz S, et al. Impact of Quorum Sensing Inhibitors on Population Dynamics of Oral Bacteria. Journal of Biology, Agriculture and Healthcare. 2019.

112. Parga A, Balboa S, Otero-Casal P, et al. New Preventive Strategy against Oral Biofilm Formation in Caries-Active Children: An In Vitro Study. Antibiotics (Basel). 2023 Jul 31;12(8).

113. Muras A, Mallo N, Otero-Casal P, et al. Quorum sensing systems as a new target to prevent biofilm-related oral diseases. Oral Dis. 2022 Mar;28(2):307–313.

114. Simon-Soro A, Mira A. Solving the etiology of dental caries. Trends in microbiology. 2015 Feb;23(2):76–82.

