## Supplementary Material for "*N*-acyl homoserine lactone signaling modulates bacterial community associated with human dental plaque"

### Table of Contents

**Page 2:** Table of contents.

**Page 3:** Fig. S1: Standard response curve of *E. coli* AHL biosensor strain JM109 pJBA132 against different concentrations of 3oC8-HSL.

**Page 4:** Fig. S2: Detection of AHLs in the spent supernatants of a dental plaque community cultured in 5% $CO_2$ atmosphere conditions.

**Page 5:** Fig. S3: AHLs were not detected in the spent supernatants of anaerobically cultured dental plaque community.

**Page 6**: Fig. S4: The Krona pie chart of the bacterial taxonomy in dental plaque community.

**Page 7:** Fig. S5: Linear discriminant analysis effect size (LEfSe) of planktonic dental plaque communities

**Page 8:** Fig. S6: Relative abundance of biofilm and planktonic microbiome associated with dental plaque.

**Page 9**: Fig. S7: Significant Operational Taxonomic Units (OTUs) in samples grown in 5% $CO_2$ atmosphere and anaerobic conditions.

**Page 10**: Table S1: AMOVA statistical tests of dental plaque communities grown in 5% $CO_2$ in both biofilm and planktonic samples.

**Page 11**: Table S2: ANOSIM statistical tests of 5% $CO_2$ dental plaque communities in both biofilm and planktonic samples.

**Page 12**: Table S3: Comparison of alpha diversity (Shannon Index) p-values between treatments in both biofilm and planktonic communities in 5% $CO_2$ condition using pairwise t-test**.**

**Page 13**: Table S4: AMOVA statistical tests of anaerobic dental plaque communities in both biofilm and planktonic samples.

**Page 14**: Table S5: ANOSIM statistical tests of anaerobic dental plaque communities in both biofilm and planktonic samples.

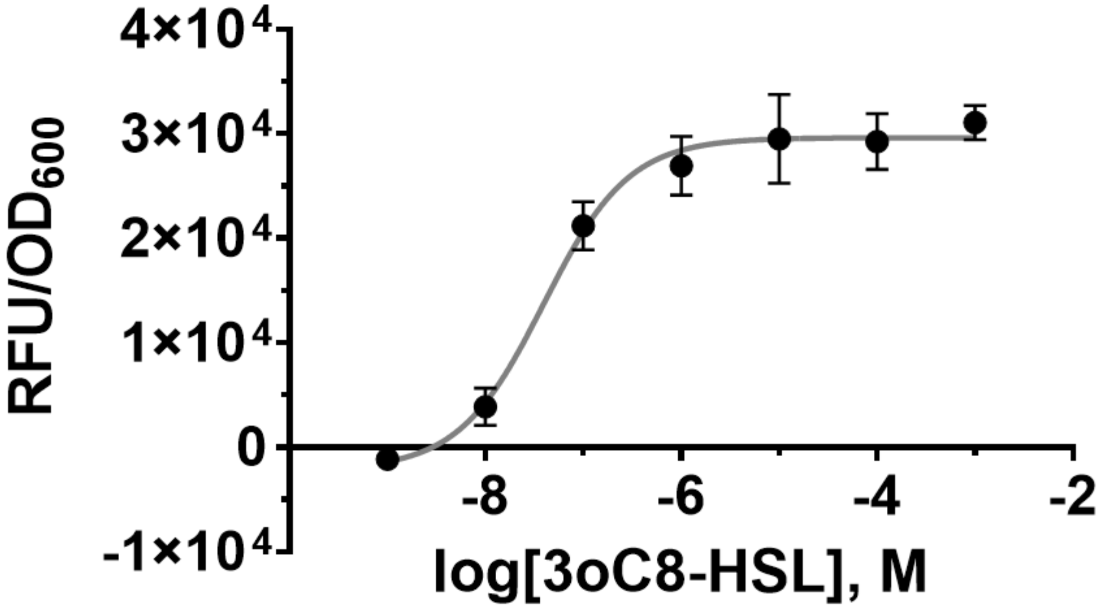


**Fig S1. Standard response curve of *E. coli* AHL biosensor strain JM109 pJBA132 against different concentrations of 3oC8-HSL.** 3oC8-HSL was freshly prepared in DMSO and diluted 1:100 into exponentially growing biosensor cultures in LB medium at indicated final concentrations. The resulting fluorescence is indicated as relative fluorescence units (RFU) per unit $OD_{600}$ of biosensor cultures. The mean and standard deviation of $RFU/OD_{600}$ values for indicated 3oC8-HSL concentrations are shown. The three-parameter nonlinear fit of $log_{10}$(3oC8-HSL concentration) versus response ($RFU/OD_{600}$) is produced by GraphPad Prism with an $EC_{50}$ of 39.42 nM and $R^2$ of 0.9611.

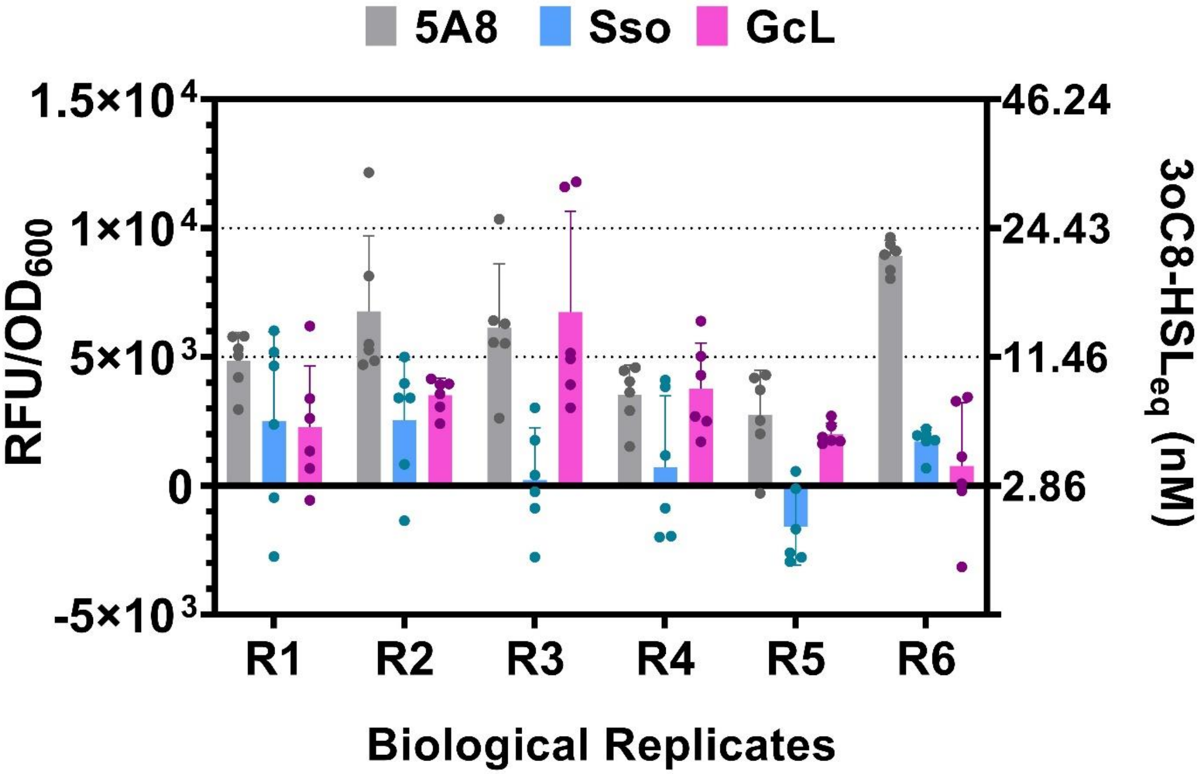


**Fig S2. Detection of AHLs in the spent supernatants of a dental plaque community cultured in 5% CO2 atmosphere conditions.** Cell-free culture supernatants of 6 biological replicates (R1 - R6) of aerobically cultured dental plaque community treated with lactonases 5A8 (inactive lactonase; control), SsoPox or GcL were incubated with *E. coli* AHL biosensor strain JM109 pJBA132 and the resulting fluorescence is indicated as relative fluorescence units (RFU) per unit $OD_{600}$ of biosensor cultures on the left Y-axis. The equivalent 3oC8-HSL concentration for the corresponding RFU/$OD_{600}$ values as shown on the right Y-axis was interpolated from the standard curve of the biosensor response against 3oC8-HSL in Fig. S1. The mean and standard deviation of RFU/$OD_{600}$ values of culture supernatants of R1 – R6 incubated with 6 biological replicates of biosensor cultures are shown after the background (mean RFU/$OD_{600}$ of 6 replicates of biosensor culture containing sterile modified Shi medium) was subtracted and the resulting values were adjusted to factor the dilution of supernatants of R1 – R6 into biosensor cultures.

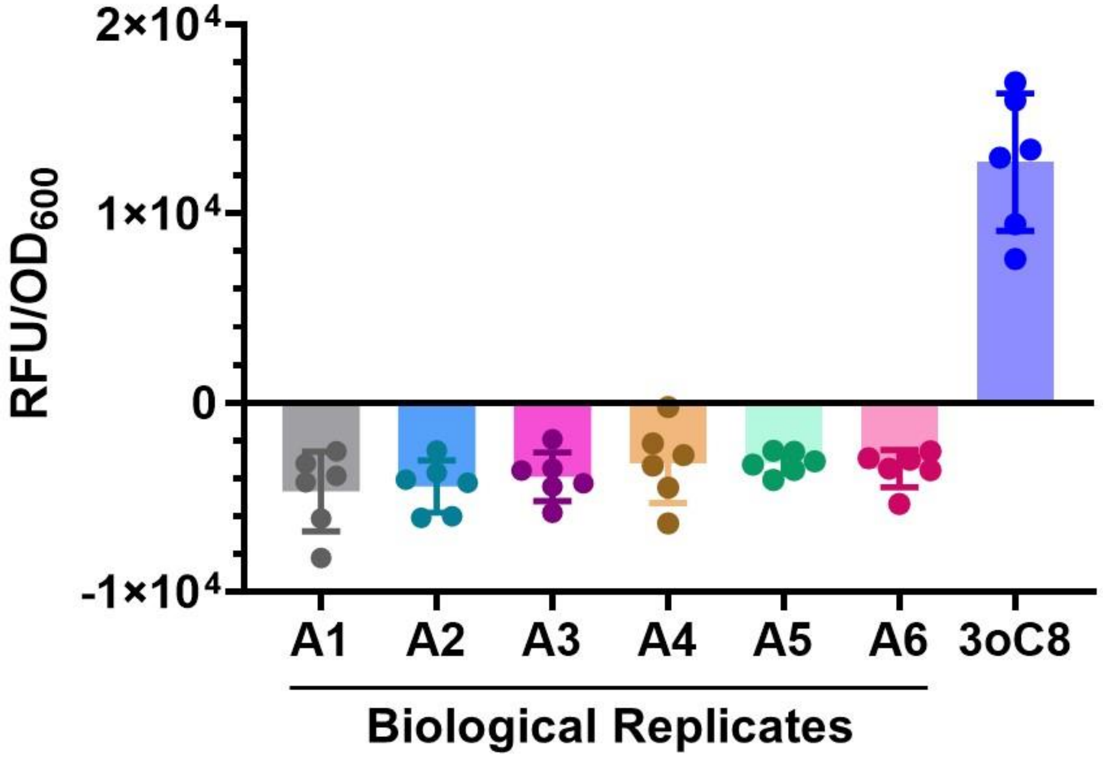


**Fig. S3. AHLs were not detected in the spent supernatants of anaerobically cultured dental plaque community.** Cell-free culture supernatants of anaerobically cultured dental plaque community (6 biological replicates A1 – A6) were incubated with *E. coli* AHL biosensor strain JM109 pJBA132 and the resulting fluorescence is indicated as relative fluorescence units (RFU) per unit $OD_{600}$ of biosensor cultures. The mean and standard deviation of RFU/$OD_{600}$ values of cell-free culture supernatants of A1 – A6 incubated with 6 biological replicates of biosensor cultures are shown after the background (mean RFU/$OD_{600}$ of 6 replicates of biosensor culture containing sterile modified Shi medium; control) was subtracted. Because the RFU/$OD_{600}$ values, from which the background signal was subtracted, were negative for A1 – A6, they were not further adjusted for dilution of supernatants into biosensor cultures. As a positive control, 20 nM 3oC8 was added to 6 replicates of biosensor culture containing sterile modified Shi medium.

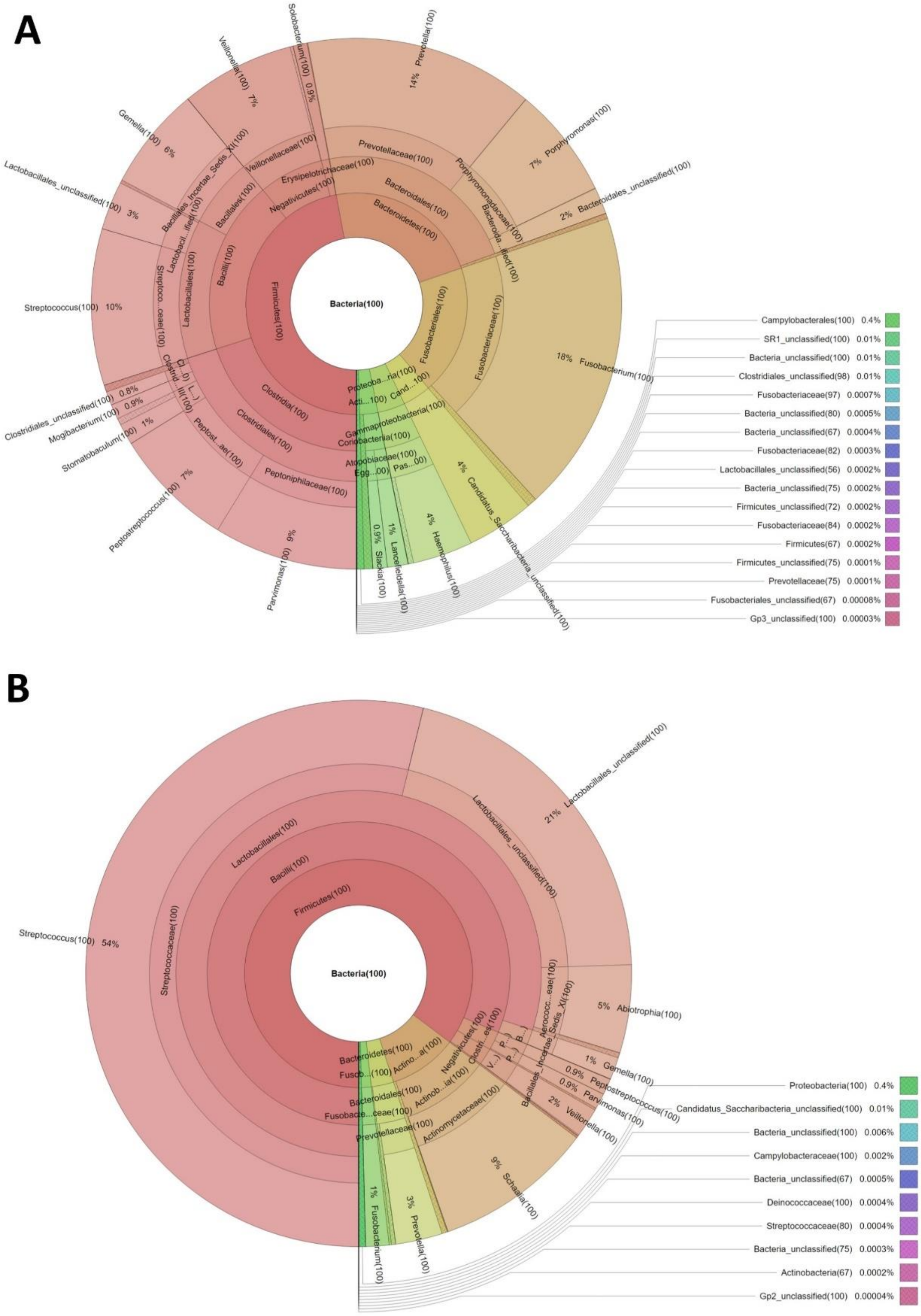


**Fig. S4: The Krona pie chart of the bacterial taxonomy in dental plaque community.** Bacteria identified from Mothur analysis in **(A)** anaerobic and **(B)** 5% $CO_2$ communities were visualized using Krona tools (https://github.com/marbl/Krona/wiki).

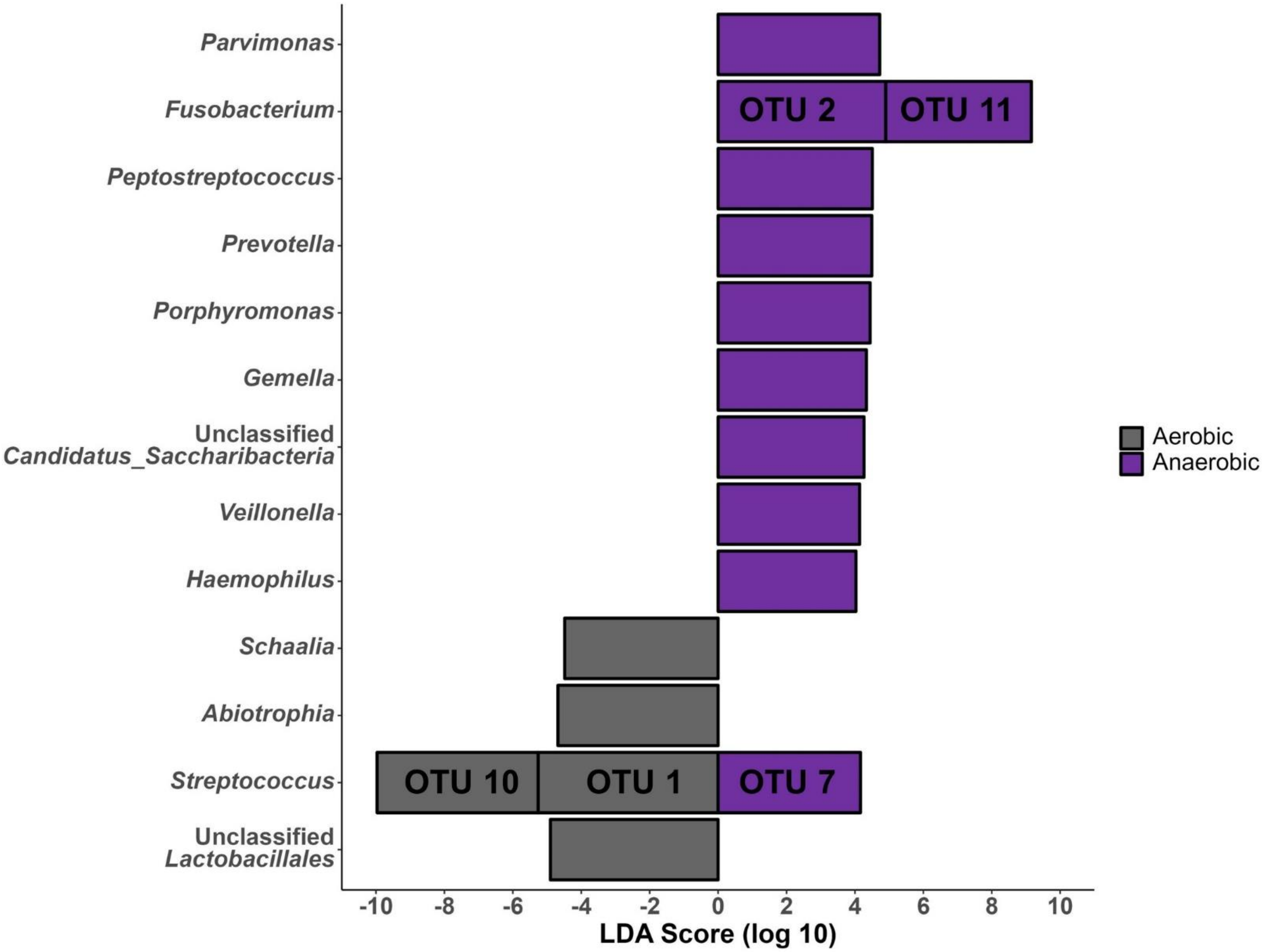


**Fig. S5: Linear discrimination analysis effect size (LEfSe) of planktonic dental plaque communities**. The bar graph of LDA scores shows the taxa with statistical difference between 5% $CO_2$ and anaerobic communities. Only taxa meeting a LDA significant threshold > 4 are shown. **Note:** *Schaalia* was formerly known as *Actinomyces*.

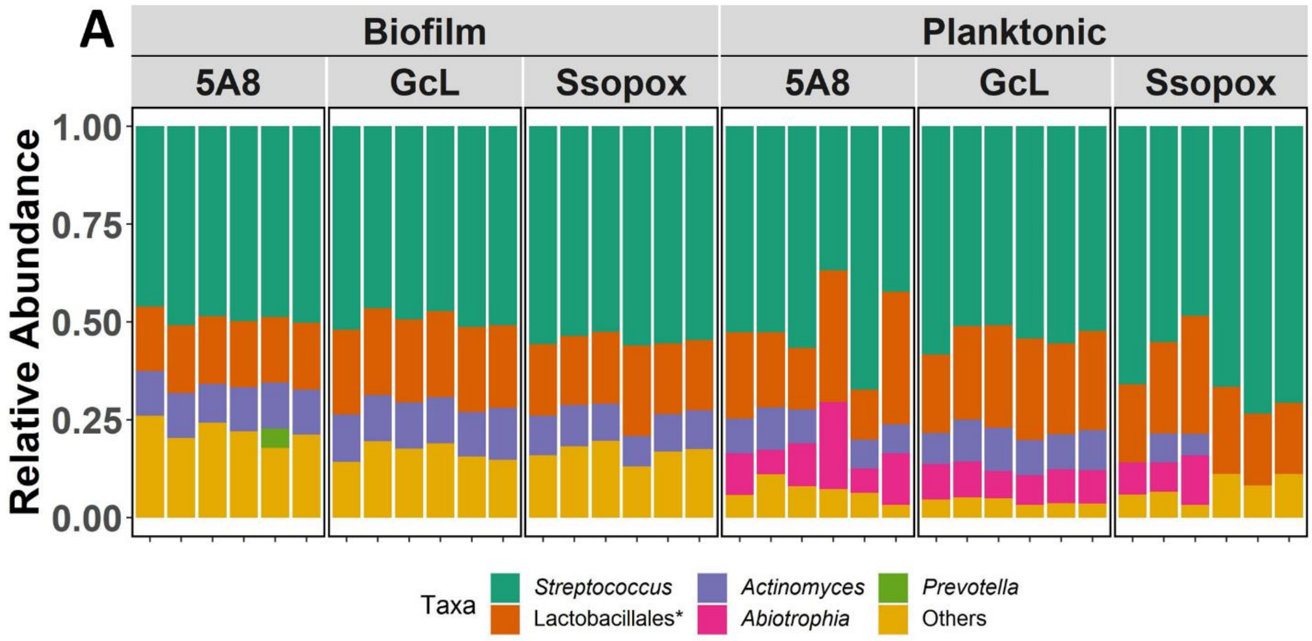


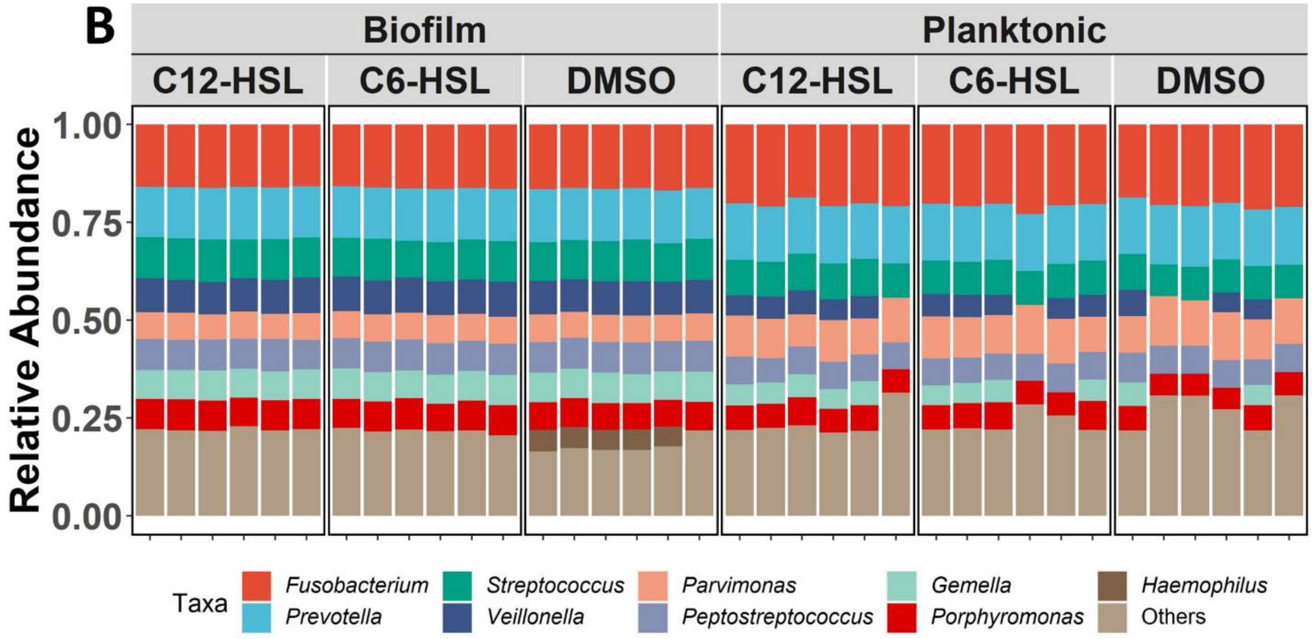


**Fig. S6. Relative abundance of biofilm and planktonic microbiome associated with dental plaque**. Taxa summary of **(A)** 5% $CO_2$ and **(B)** anaerobic conditions per sample.

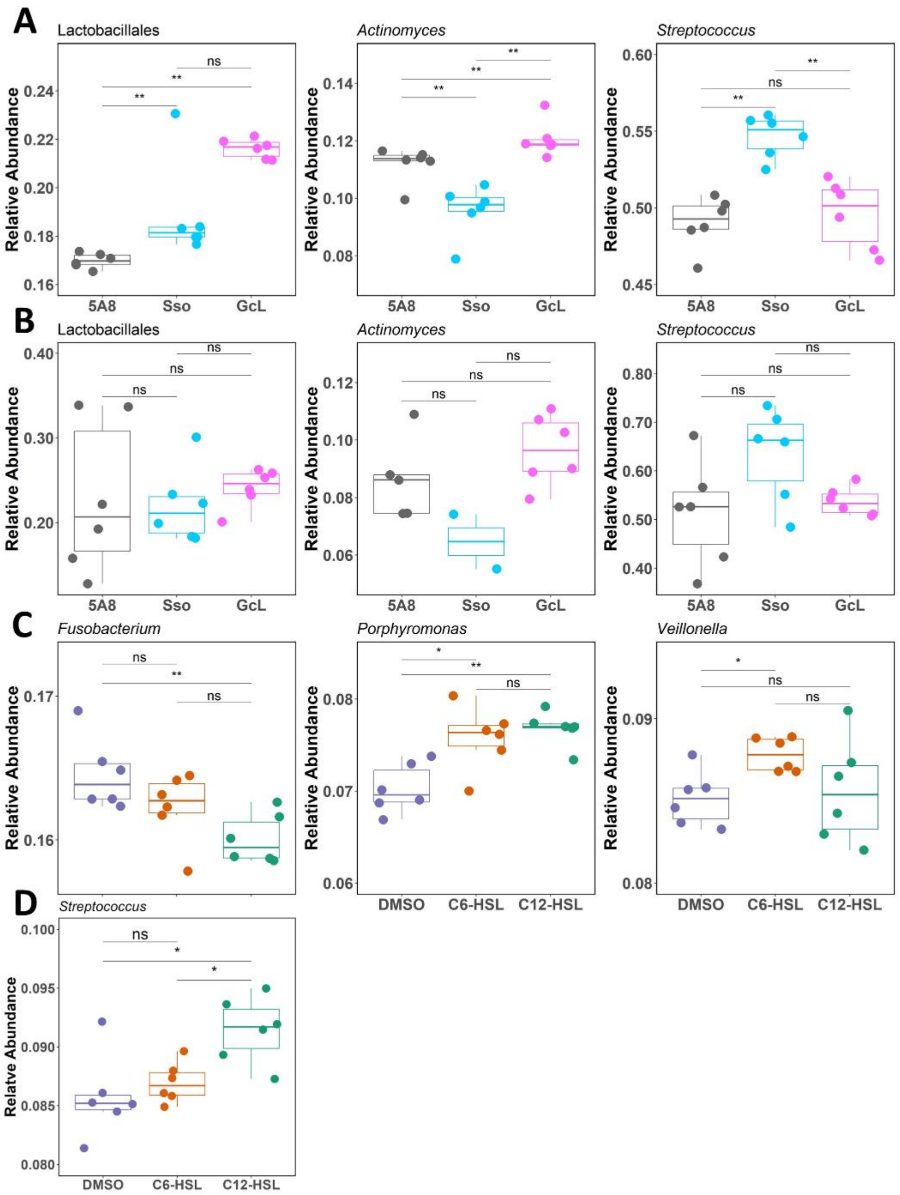


**Fig. S7: Significant Operational Taxonomic Units (OTUs) in samples grown in 5% CO2 atmosphere and anaerobic conditions.** Samples are colored by treatment (gray: control (5A8); blue: SsoPox; pink: GcL). **(A)** Major Biofilm and **(B)** planktonic taxa difference in ***5% CO₂ condition***. **Note:** There were no significant differences in major bacterial species between the treatments in the planktonic communities. **(C)** Biofilm and **(D)** planktonic bacterial species with significant differences in ***anaerobic conditions*.** Differences in relative abundance by treatments were tested for by using a Wilcoxon test, and significance values are indicated as - ****$p < 0.0005$*, **$p < 0.005$ and *$p < 0.05$, ns $p > 0.05$.*

**Table S1: AMOVA statistical tests of 5% $CO_2$ dental plaque communities in both biofilm and planktonic samples.**

| Samples Comparison | Statistical factor | Among | Within | Total | P-values |
|---|---|---|---|---|---|
| **Biofilm** | | | | | |
| 5A8-GcL-SsoPox | SS | 0.054964 | 0.035265 | 0.0902414 | < 0.001* |
| | Df | 2 | 15 | 17 | |
| | MS | 0.0274882 | 0.002351 | | |
| | Fs | 11.6921 | | | |
| 5A8-GcL | SS | 0.0302529 | 0.0203439 | 0.0505968 | 0.002* |
| | Df | 1 | 10 | 11 | |
| | MS | 0.0302529 | 0.00203439 | | |
| | Fs | 14.8708 | | | |
| 5A8-SsoPox | SS | 0.0256429 | 0.0248728 | 0.0505157 | 0.002* |
| | Df | 1 | 10 | 11 | |
| | MS | 0.0256429 | 0.00248728 | | |
| | Fs | 10.3096 | | | |
| GcL-SsoPox | SS | 0.0265687 | 0.0253134 | 0.0518821 | < 0.001* |
| | df | 1 | 10 | 11 | |
| | MS | 0.0265687 | 0.00253134 | | |
| | Fs | 10.4959 | | | |
| **Planktonic** | | | | | |
| 5A8-GcL-SsoPox | SS | 0.0779351 | 0.227876 | 0.305811 | 0.047* |
| | Df | 2 | 15 | 17 | |
| | MS | 0.0389675 | 0.0151917 | | |
| | Fs | 2.56505 | | | |
| 5A8-GcL | SS | 0.0172218 | 0.160015 | 0.177237 | 0.371 |
| | Df | 1 | 10 | 11 | |
| | MS | 0.0172218 | 0.0160015 | | |
| | Fs | 1.07626 | | | |
| 5A8-SsoPox | SS | 0.0454945 | 0.205831 | 0.251325 | 0.143 |
| | Df | 1 | 10 | 11 | |
| | MS | 0.0454945 | 0.0205831 | | |
| | Fs | 2.21029 | | | |
| GcL-SsoPox | SS | 0.0541863 | 0.0899062 | 0.144092 | 0.018* |
| | df | 1 | 10 | 11 | |
| | MS | 0.0541863 | 0.00899062 | | |
| | Fs | 6.026299 | | | |

**Table S2: ANOSIM statistical tests of 5% $CO_2$ dental plaque communities in both biofilm and planktonic samples.**

| Biofilm | | |
|---|---|---|
| **Sample Comparison** | **R-values** | **P-values** |
| 5A8-GcL-SsoPox | 0.857202 | < 0.001* |
| 5A8-GcL | 1 | 0.002* |
| 5A8-SsoPox | 0.733333 | 0.002* |
| GcL-SsoPox | 0.831481 | < 0.001* |
| **Planktonic** | | |
| **Sample Comparison** | **R-values** | **P-values** |
| 5A8-GcL-SsoPox | 0.304733 | 0.003* |
| 5A8-GcL | 0.255556 | 0.007* |
| 5A8-SsoPox | 0.190741 | 0.081 |
| GcL-SsoPox | 0.514815 | 0.01* |

**Table S3: Comparison of alpha diversity (Shannon Index) p-values between treatments in both biofilm and planktonic communities in 5% $CO_2$ condition using pairwise t-test.**

| **Planktonic** | | | |
|---|---|---|---|
| | Control | GcL | SsoPox |
| Control | - | 0.74 | 0.025 |
| GcL | - | - | 0.031 |
| **Biofilm** | | | |
| | Control | GcL | SsoPox |
| Control | - | 0.013 | 0.00083 |
| GcL | - | - | 0.099 |

**Table S4: AMOVA statistical tests of anaerobic dental plaque communities in both biofilm and planktonic samples.**

| Biofilm | | | | | |
|---|---|---|---|---|---|
| **Samples Comparison** | **Statistical factor** | **Among** | **Within** | **Total** | **P-values** |
| DMSO-C12-HSL-C6-HSL | SS | 0.00392449 | 0.0114633 | 0.0153878 | < 0.001* |
| | Df | 2 | 15 | 17 | |
| | MS | 0.00196224 | 0.000764222 | | |
| | Fs | 2.56764 | | | |
| C12-HSL-C6-HSL | SS | 0.00145441 | 0.00767769 | 0.00913209 | 0.004* |
| | Df | 1 | 10 | 11 | |
| | MS | 0.00145441 | 0.000767769 | | |
| | Fs | 1.89433 | | | |
| DMSO-C12-HSL | SS | 0.00224788 | 0.00739184 | 0.00963972 | 0.002* |
| | Df | 1 | 10 | 11 | |
| | MS | 0.00224788 | 0.000739184 | | |
| | Fs | 3.04102 | | | |
| DMSO-C6-HSL | SS | 0.00218445 | 0.00785712 | 0.0100416 | 0.001* |
| | df | 1 | 10 | 11 | |
| | MS | 0.00218445 | 0.000785712 | | |
| | Fs | 2.78022 | | | |
| **Planktonic** | | | | | |
| DMSO-C12-HSL-C6-HSL | SS | 0.00940708 | 0.043356 | 0.0527631 | 0.126 |
| | Df | 2 | 15 | 17 | |
| | MS | 0.00470354 | 0.0028904 | | |
| | Fs | 1.6273 | | | |

**Table S5: ANOSIM statistical tests of anaerobic dental plaque communities in both biofilm and planktonic samples.**

| Biofilm | | |
|---|---|---|
| **Sample Comparison** | **R-values** | **P-values** |
| DMSO-C12-HSL-C6-HSL | 0.541152 | < 0.001* |
| C12-HSL-C6-HSL | 0.303704 | 0.007* |
| DMSO-C12-HSL | 0.733333 | 0.004* |
| DMSO-C6-HSL | 0.62037 | 0.002* |
| **Planktonic** | | |
| **Sample Comparison** | **R-values** | **P-values** |
| DMSO-C12-HSL-C6-HSL | 0.138272 | 0.071 |
